# TRF1 prevents permissive DNA damage response, recombination and Break Induced Replication at telomeres

**DOI:** 10.1101/697979

**Authors:** Rosa Maria Porreca, Pui Pik Law, Emilia Herrera-Moyano, Roser Gonzalez-Franco, Alex Montoya, Peter Faull, Holger Kramer, Jean-Baptiste Vannier

## Abstract

Telomeres are a significant challenge to DNA replication and are prone to replication stress and telomere fragility. The shelterin component TRF1 facilitates telomere replication but the molecular mechanism remains uncertain. By interrogating the proteomic composition of telomeres, we show that telomeres lacking TRF1 undergo protein composition reorganisation associated with a DNA damage response and chromatin remodelers. Surprisingly, TRF1 suppresses the accumulation of promyelocytic leukemia (PML) protein, BRCA1 and the SMC5/6 complex at telomeres, which is associated with increased Homologous Recombination (HR) and TERRA transcription. We uncovered a previously unappreciated role for TRF1 in the suppression of telomere recombination, dependent on SMC5 and also POLD3 dependent Break Induced Replication at telomeres. We propose that TRF1 facilitates S-phase telomeric DNA synthesis to prevent illegitimate mitotic DNA recombination and chromatin rearrangement.

## Introduction

Telomeres are specialised nucleoprotein structures at the ends of chromosomes, composed of repetitive sequences (TTAGGG repeats in mammals) (Moyzis et al., 1988), long non-coding RNA called TERRA and six associated proteins, TRF1, TRF2, POT1a/b, RAP1 and TIN2, that form the shelterin complex (de Lange, 2005). These capping structures have the crucial function of maintaining genome stability by protecting the chromosome end from being recognised as DNA double strand breaks (DSBs) (Palm & de Lange, 2008). They also represent challenging structures for the replication machinery, which is associated to telomere fragile sites (Martinez et al., 2009; McNees et al., 2010; Sfeir et al., 2009; Vannier, Pavicic-Kaltenbrunner, Petalcorin, Ding, & Boulton, 2012). Telomere fragility is identified by the formation of multitelomeric signals (MTS), where telomeres appear as broken or decondensed, resembling the common fragile sites (CFS) observed at non telomeric loci after treatment with aphidicolin (APH). TRF1 facilitates the progression of the replication fork at telomeres, by recruiting specialised DNA helicase BLM, which in turn resolve secondary structures, similar to fission yeast ortholog Taz1 (Lee, Arora, Wischnewski, & Azzalin, 2018; Martinez et al., 2009; Miller, Rog, & Cooper, 2006; Sfeir et al., 2009).

During tumorigenesis, cancer cells can achieve replicative immortality by activation of telomere maintenance mechanisms. The majority of cancer cells reactivate telomerase, while a minority (10-15%) uses an alternative mechanism named ALT for alternative lengthening of telomeres (Bryan, Englezou, Dalla-Pozza, Dunham, & Reddel, 1997; Kim et al., 1994). Intriguingly, ALT is characterised by the appearance of ALT-associated PML bodies (APBs), specialised sites where a subset of telomeres co-localises with PML protein and several DNA repair and homologous recombination (HR) proteins (Draskovic et al., 2009; G. Wu, Lee, & Chen, 2000; Yeager et al., 1999). ALT telomeres can be maintained by more than one mechanism of recombination. Indeed, in yeast, two different ALT-like pathways have been described: Type I, requires Rad51 to mediate the invasion of a homologous sequence, while Type II is Rad51 independent and rely on Rad52 dependent elongation mechanism, which consists in the annealing of ssDNA regions. Both Type I and II mechanisms require the DNA polymerase Pol32, which initiates DNA synthesis for several kilobases, in a process known as Break Induced Replication (BIR) (Ira & Haber, 2002). Recently, multiple groups have revisited this Rad51 independent DNA synthesis repair pathway at mammalian ALT telomeres (Dilley et al., 2016; Garcia-Exposito et al., 2016; Roumelioti et al., 2016). Mammalian BIR is dependent on POLD3 and POLD4, subunits of DNA polymerase delta and orthologs of yeast Pol32. ALT cells present increased DNA damage response (DDR) and several studies have underlined the contribution of replication stress to ALT-mediated telomere extension (Arora et al., 2014; K. E. Cox, Marechal, & Flynn, 2016; Pan et al., 2017). However, the molecular mechanisms initiating recombination in ALT cells are still unclear.

In order to gain insight into the chromatin composition of telomeres undergoing replication stress, we performed Proteomics of Isolated Chromatin segments (PICh), using *TRF1* conditional knock-out Mouse Embryonic Fibroblasts (MEFs, telomerase positive). Surprisingly, we found that telomeres lacking TRF1 are enriched in SMC5/6, DNA polymerase δ (POLD3), and chromatin remodeling factors known to be associated with ALT telomeres. These cells also present additional DNA damage and recombination hallmarks such as formation of APBs, mitotic DNA synthesis at telomeres, a feature of BIR, recruitment of chromatin remodeling factors and increased TERRA levels. Further investigation using specific shRNAs against the SMC5/6 complex or POLD3 revealed how these two complexes are key regulators of the recombination signature identified in *TRF1* deleted cells. Taken together, these results strongly identify TRF1 as a central player in preserving telomeric chromatin against HR, induced by DNA replication stress, and particularly POLD3 dependent-mitotic DNA synthesis.

## Results

### Capture of TRF1 depleted telomeres by PICh reveals drastic changes in the chromatin composition

To isolate and identify the chromatin composition of TRF1 depleted telomeres, we employed Proteomics of Isolated Chromatin segments (PICh), a powerful and unbiased technique that uses a desthiobiotinylated oligonucleotide complementary to telomeric repeat sequences to specifically pull down telomeric chromatin (Dejardin & Kingston, 2009). We performed PICh in MEFs harboring a *TRF1* conditional allele. MEFs lacking *TRF1* are well known to undergo replicative stress; however, they can grow for up to 8 days before entering senescence, making them optimal for investigating replication stress at telomeres (Martinez et al., 2009; Sfeir et al., 2009). Cells were transduced twice (day 0 and 3) with a CRE or GFP control adenovirus and collected 7 days after the first transduction, as indicated in the timeline (Figure 1A). Excision of exon 1 of *TRF1* by CRE recombinase (Sfeir et al., 2009) resulted in the expected loss of TRF1 protein as determined by immunoblotting (Figure 1B). Cells were fixed and isolation of telomeres was performed using a probe complementary to TTAGGG repeats or a scrambled probe as a negative control. Finally, telomeric chromatin was isolated from both control cells (wt) and *TRF1* deleted cells before mass spectrometry identification (Figure 1C). We identified a list of 1306 proteins that was subjected to refinement in order to remove unspecific bound proteins or contaminants found with the scrambled probe (see experimental procedure for detailed description). Based on the analysis of *label free* quantification (LFQ intensities), we found 119 proteins presenting a gain of abundance at TRF1 depleted telomeres (Log2>-2) and 206 factors were displaced from these telomeres (Log2>2), considering that a cut-off for differential expression is set to log2 fold change (TRF1deletion/wt)> |2| and -Log (p-value) >1 (Figure 1D). Amongst these 206 proteins, we found TRF1, as expected due to the knock-out of its gene, but also one component of the CST complex (CTC1), important player in the efficient restart of stalled replication forks at telomeres (Gu et al., 2012) and recruited through POT1b interaction (P. Wu, Takai, & de Lange, 2012). Interestingly, POT1b is also less abundant at TRF1 depleted telomeres (Figure 1D-E). On the other end, the group of 119 proteins enriched in *TRF1* deleted cells includes several factors involved in structural maintenance of chromosomes (SMC), HR and DNA damage response (Figure 1D-E-F), such as the MRN complex (MRE11, RAD50 and NBS1). The identification of 53BP1 recruited to TRF1 depleted telomeres (Figure 1D-F) acts as a positive marker for the specificity of this proteomic analysis, as reported before in (Martinez et al., 2009; Sfeir et al., 2009). Moreover, we could identify drastic and previously uncharacterised changes of the telomeric proteome at telomeres undergoing replication stress presented hereafter.

**Figure 1.**
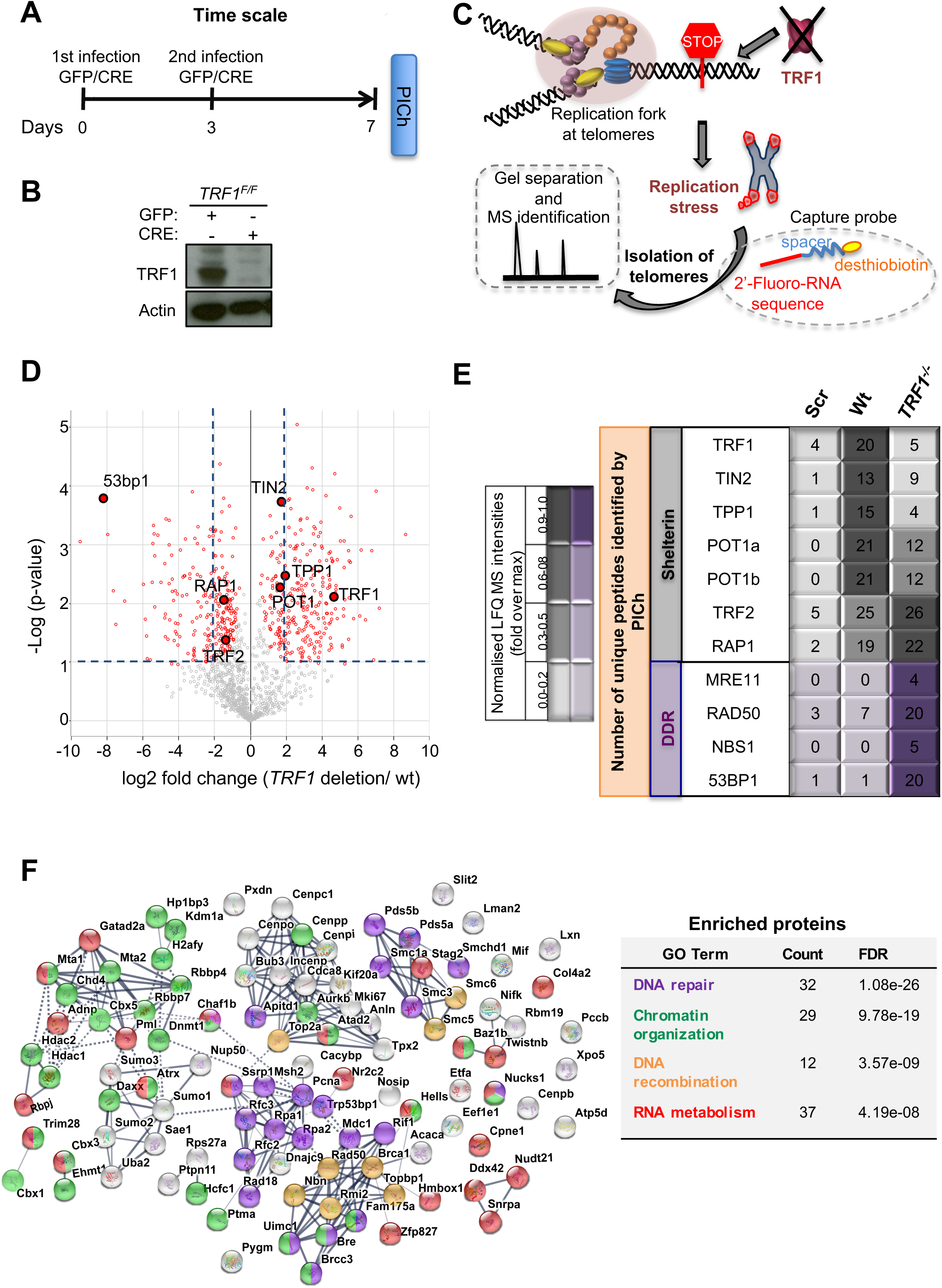
Proteomics of isolated chromatin segments (PICh) of TRF1 depleted mouse telomeres. (A) Overview of experimental timeline aimed at performing PICh experiment after induction of *TRF1* deletion. *TRF1^F/F^* MEFs were infected twice (day 0 and 3) with adenovirus containing either GFP-control or CRE and collected at day 7 for PICH experiments. **(B)** Western blot showing deletion of *TRF1* in MEFs after infections with CRE Adenovirus, at day 7 as in A. **(C)** Schematic representation of the PICh analysis performed to detect chromatin changes occurring at telomeres upon *TRF1* deletion. **(D)** Volcano Plot based on LFQ intensities of proteins. Cut off for differential expression were set to log2 fold change (TRF1deletion/wt)> |2| and -Log (p-value) >1. **E)** Table listing shelterin components and some of the DNA damage response (DDR) factors identified. The corresponding number of unique peptide isolated is indicated for each factor of interest. Relative LFQ intensity abundance profiles were visualised in the form of a heat-map, by scaling each protein intensity to the maximum intensity across conditions. Light to darker colors indicate increasing relative protein abundance. **(F)** Connectivity map for proteins recruited at telomeres upon TRF1 deletion using string-db.org software. Solid lines, represents strong direct interactions, while dashed lines represent no evidence for direct interaction. In violet, DNA damage and repair proteins; in orange, factors belonging to DNA repair specifically involved in DNA recombination process; while in green and red, important factors for chromosome maintenance and factors involved in RNA metabolism, respectively. Source data are provided as a Source Data File.

### TRF1 suppresses APBs formation and HR at telomeres

Interestingly, *TRF1* deficient MEFs present a telomeric enrichment for factors involved in HR and chromatin remodeling (NurD complex, BRCA1, SMC5/6 and PML) that are usually abundant at ALT telomeres (Figure 1D-2A) (Conomos, Reddel, & Pickett, 2014; Draskovic et al., 2009; Marzec et al., 2015; Potts & Yu, 2007). To validate the specific association of some of these factors with TRF1 depleted telomeres in telomerase positive MEFs, we carried out chromatin immunoprecipitation (ChIP) experiments using ChIP-grade specific antibodies followed by telomeric dot-blot. TRF1 antibody was used as a negative control for our experiment, while the recruitment of BRCA1, BAZ1b, and some subunits of the nucleosome remodeling and deacetylase (NurD) complex (p66a, MTA1, ChD4, zinc-finger protein ZNF827) was assessed. For all these factors, with the exception of p66a for which no statistical significance was achieved, we observed a specific enrichment at telomeres upon *TRF1* deletion (Figure 2B; Figure S1A-B). In addition, to confirm the presence of PML at replication stress induced telomeres, as suggested by our PICh data (Figure 2A), we performed immuno-FISH and scored for the formation of APBs. We observed a two-fold increase in the number of co-localisations between PML and telomeres in *TRF1^-/-^* MEFs compared to control cells (Figure 2C). Overall these data demonstrate that telomeres undergoing replication stress favor the recruitment of chromatin remodeler, HR factors and the formation of APBs, considered a platform of recombination for chromosome ends (Cesare & Reddel, 2010). This suggests a role of TRF1 in suppressing recombination events as well as many other phenotypic features related to ALT. Hence, to test this hypothesis, we revisited the incidence of telomeric sister chromatid exchanges (T-SCE) using chromosome orientation FISH (CO-FISH) in *TRF1* deficient cells (Figure 2D). We identified an increase in T-SCE in *TRF1^-/-^* MEFs (2.8%) compared to control cells (0.4%) (Figure 2D). This result is at odds with previous publications where T-SCE events detected at TRF1 depleted telomeres were not significantly enriched, with only 1% of T-SCEs detected compared to 0.1% in wt cells (Martinez et al., 2009; Sfeir et al., 2009). In fact, this discrepancy might be explained by the difference in timing for the analysis of T-SCEs in *TRF1* deficient cells. Both publications report the lack of recombination effect by T-SCEs at 3 or 4 days after *TRF1* loss, while we generally carry our investigations at day 7. Therefore, we repeated the experiments in *TRF1^-/-^* cells at different time points post infection: day 4 and day 7, finding respectively 1.6% and 2.8% of T-SCEs per chromosome end (Figure S2, left graph), indicating a lower % of T-SCE events happening at earlier time point. A second distinct difference with previous reports is the type of telomere signal exchanges that we analysed. As in Sfeir et al., 2009, all types of telomere signal exchanges (e.g. the exchanges appearing at single chromatids and the reciprocal exchanges at both chromatids) were considered. However, Martinez et al., 2009 only refers to reciprocal exchanges at both chromatids. Thus, we next classified T-SCEs detected in *TRF1* deficient MEFs into these two different types (single and double) and found that 4 days post infection only T-SCEs at single chromatids were significantly increased (Figure S2, right graph), while the reciprocal exchanges were not enhanced at TRF1 depleted telomeres (Figure S2, middle graph). Therefore, our detailed analysis of the nature and timing of T-SCEs in TRF1 deficient MEFs is in line with the previous literature. Moreover, it demonstrates the unappreciated role of TRF1 in suppressing HR and suggests that the initial recombination events happening at replication stressed telomeres could be generated by the BIR pathway (single chromatid exchanges) (Roumelioti et al., 2016).

**Figure 2.**
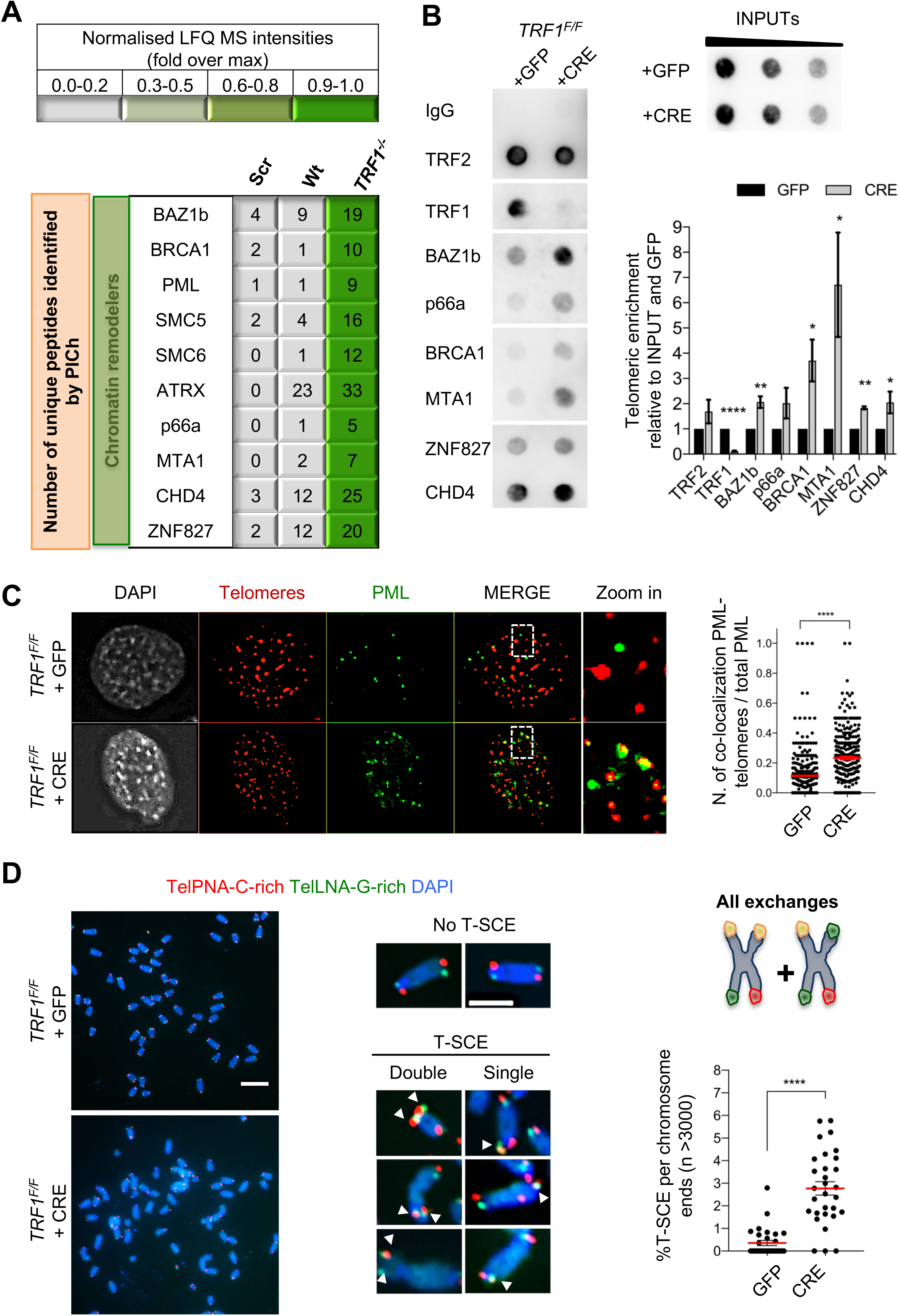
Recombination factors are recruited at TRF1 depleted telomeres. **(A)** Table listing chromatin remodelers identified. The corresponding number of unique peptide isolated is indicated for each factor of interest. Same as in Figure 1, light to darker colors indicate increasing relative protein abundance. **(B**) Validation of chromatin remodeler factors by ChIP-dot blot analysis in wt (+GFP) and *TRF1^-/-^* (+CRE) conditions using ChIP grade antibodies against chosen factors after chromatin preparation from MEFs. The blot was revealed with a DIG-Tel-C-rich probe. ChIP signals were normalised to DNA input and GFP control. Data are represented as telomeric enrichment of proteins relative to GFP (n=3) ± SEM. P values, two-tailed student t-test (*, P < 0.05;**, P < 0.01; ****, P < 0.0001). (**C)** Representative image of Immunofluorescence showing co-localisation of Telomeres (red) with PML (green) in MEFs nuclei (DAPI) treated with GFP and CRE. Data are represented as number of Telomeres-PML co-localising foci divided by the total number of PML present per nucleus (n=300 nuclei) and are shown as mean (red line) ± SEM. P values, two-tailed student t-test (****, *P* < 0.0001). Source data are provided as a Source Data File. **(D)** Representative images of the chromosome oriented CO-FISH assay <u>with denaturation</u>, used to score for telomeric T-SCEs in *TRF1^F/F^* MEFs infected with GFP or CRE. Telomeres are labeled with TelPNA-C-rich-Cy3 (red) and TelLNA-G-rich-FAM (green), while chromosomes are counterstained with DAPI (blue). Scale bar, 10 µm. Enlarged intersections show the difference between a chromosome with No T-SCE (top) and a chromosome with T-SCE (bottom). T-SCE images show double T-SCEs (left) and single chromatid events (right). Scale bar, 2 µm. For quantification, T-SCE was considered positive when involved in a reciprocal exchange of telomere signal with its sister chromatid (both telomeres yellow) and for asymmetrical exchanges at single chromatid (one telomere yellow). Data are indicated as % of T-SCE per sister telomere. The mean values (n=>3000 chromsome ends) ± SEM are indicated. P value, two-tailed student t-test (****, *P* < 0.0001).

### TRF1 depletion causes TERRAs upregulation

Since depleting telomeres of TRF1 induces the formation of APBs and the increase of HR, we next decided to revisit the role of TRF1 in telomere transcription, as TERRA molecules are proposed to regulate telomere recombination (Yu et al., 2014). Previous studies have reported *in-vivo* interactions between TRF1 and TERRA (Deng, Norseen, Wiedmer, Riethman, & Lieberman, 2009) and also a possible transcriptional regulation by TRF1 through a mechanism involving RNA polymerase II -TRF1 interaction (Schoeftner & Blasco, 2008). However, the role of TRF1 regulating telomere transcription appears complex since contrasting results have been reported by different groups in both human and mouse cell lines (Lee et al., 2018; Schoeftner & Blasco, 2008; Sfeir et al., 2009). We performed both RNA dot-blot and Northern-blot analyses showing a significant increase in TERRA molecules upon loss of *TRF1* in immortalised MEFs, 7 days after transduction (Figure 3A-B) but also at earlier time point (day 4) and in primary MEFs (Figure S3A-B-C). Collectively, we identify an increase in TERRA molecules upon TRF1 removal from telomeres, confirming transcriptional and telomeric chromatin changes in TRF1 depleted cells. Particularly, the TERRAs molecules increasing upon *TRF1* deletion have high molecular weight and can only be detected when an alkaline treatment is performed during Northern-blotting (Figure 3B; S3D). In addition, we carried out TERRA-FISH (Figure 3C), confirming a significant increase in numbers and intensity of TERRA foci per nucleus deficient for *TRF1* (Figure 3C). Taken together, these results suggest that TRF1 dependent replication stress at telomeres changes the telomeric chromatin composition by recruiting specific chromatin remodelers, which directly or indirectly affect telomere transcription and contribute to the formation of APBs, platform of recombination. The presence of these ALT-hallmarks suggests that TRF1 depleted telomeres present some similarities with ALT telomeres. However, the absence of telomere heterogeneity, c-circle formation and still presence of telomerase activity (Figure S4A-B-C) also suggest that this ALT-like phenotype is not complete.

**Figure 3.**
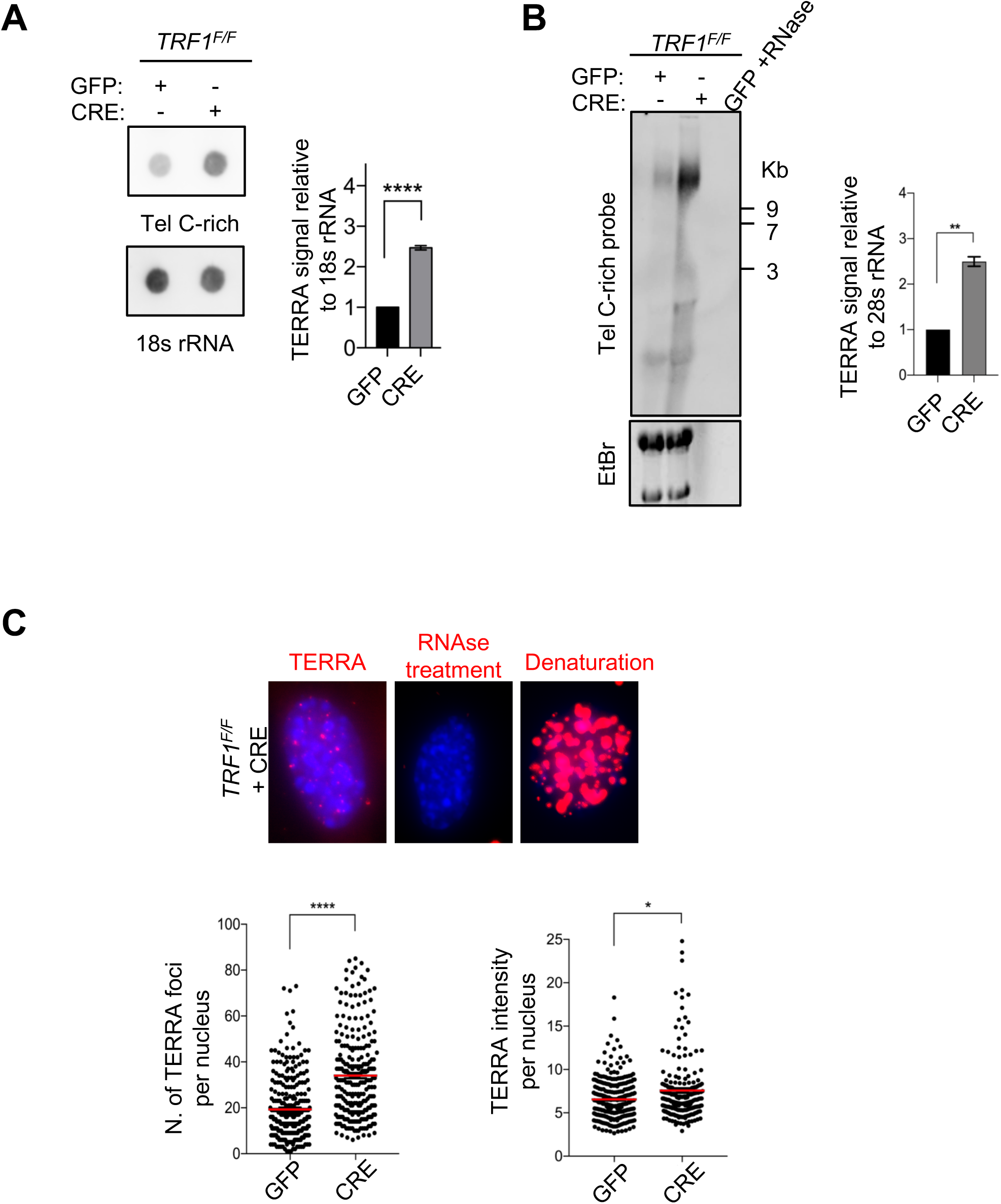
TRF1 depletion causes TERRAs upregulation. **(A)** RNA dot blot analysis in wt and *TRF1* deleted MEFs. The blot was revealed with a DIG-Tel-C-rich probe or 18s rRNA as a control. TERRA signals were normalised to 18s rRNA and GFP control (n = 3) ± SEM. P values, two-tailed student t-test (****, *P* < 0.0001). **(B)** TERRA detection by Northern blotting upon *TRF1* deletion. The blot was revealed with a DIG-Tel-C-rich probe (upper part). Ethidium bromide (EtBr) staining (bottom) of rRNAs was used as loading control. TERRA signals were normalized to 28s rRNA signal from EtBr staining (n = 2) ± SEM. P values, two-tailed student t-test (**, *P* < 0.01). **(C)** Representative images of TERRA-FISH experiment (top panel) showing the difference between cells stained with TERRA (red), negative control with RNAse A treatment and positive control after denaturation. TERRA-FISH quantification (bottom panel) in wt (+GFP) and *TRF1^-/-^* (+CRE) conditions. Graphs are representing the number of TERRA foci (left) and TERRA intensity (right) (n=250). Red lines represent mean values, two-tailed student t-test (****, P < 0.0001); Mann-Whitney test used for TERRA intensity quantification (*, P < 0.05).

### TRF1 suppresses mitotic DNA synthesis at telomeres

Since the denaturing CO-FISH experiments in *TRF1* deficient cells identified single chromatid exchanges that are proposed to be reminiscent of BIR events, an HR alternative pathway required in G2-M phase (Roumelioti et al., 2016), we tested whether TRF1 depleted telomeres trigger non S-phase DNA synthesis. We performed a pulse with 5-bromo-2-deoxyuridine (BrdU) for 2 hours (Figure 4A) before carrying out BrdU immunofluorescence at telomeres in interphase cells (Figure 4B-C). Only non S-phase cells were counted in this experiment, based on the formation of clear BrdU foci (Dilley et al., 2016; Nakamura, Morita, & Sato, 1986) (Figure 4B). *TRF1^-/-^* MEFs display elevated BrdU incorporation at telomeres, showing eight times more telomere synthesis (positive cells with more than 5 foci) compared to control cells (Figure 4C). To investigate DNA synthesis happening exclusively in mitosis, so-called MiDAS (Minocherhomji et al., 2015), we performed a similar experiment in metaphases. After incubating wt and *TRF1* deficient MEFs with 5-ethynyl-2-deoxyuridine (EdU) and colcemid for 1-hour, mitotic cells were collected to analyse EdU incorporation on metaphase chromosomes (Figure 4A). We scored for telomeric and non-telomeric EdU foci (mitotic DNA synthesis) and found that CRE induced cells had a significant increase in telomeric mitotic DNA synthesis compared to the GFP control cells (Figure 4D). This result confirms that TRF1 depleted telomeres present an increased level of non-S-phase DNA synthesis, similar to what is observed in ALT cells. In addition, analysis of EdU incorporation in metaphase spreads allowed us to distinguish between conservative BIR associated DNA synthesis and HR semi-conservative DNA synthesis (Min, Wright, & Shay, 2017). In the first case, EdU would be labeled on a single chromatid (Figure 4E, upper panel), while in the latter, EdU would localise to both chromatids (Figure 4E, bottom panel). Thus, to assess the mechanism of DNA synthesis in *TRF1* deleted cells, the pattern of EdU incorporation on metaphase chromosomes was further investigated (Figure 4F). Non-telomeric (upper panel) and telomeric (middle panel) EdU foci formed mainly on a single chromatid. In fact, 72% of the mitotic DNA synthesis at non-telomeric sites localised to a single chromatid, while the remaining 28% of the signal was present at both chromatids (Figure 4F, upper panel). This result is even more striking when EdU signal was restricted to telomeres, with almost all the co-localisation being present at single chromatids (95%). These observations suggest that TRF1 is crucial for the suppression of mitotic DNA synthesis mediated by BIR at telomeres.

**Figure 4.**
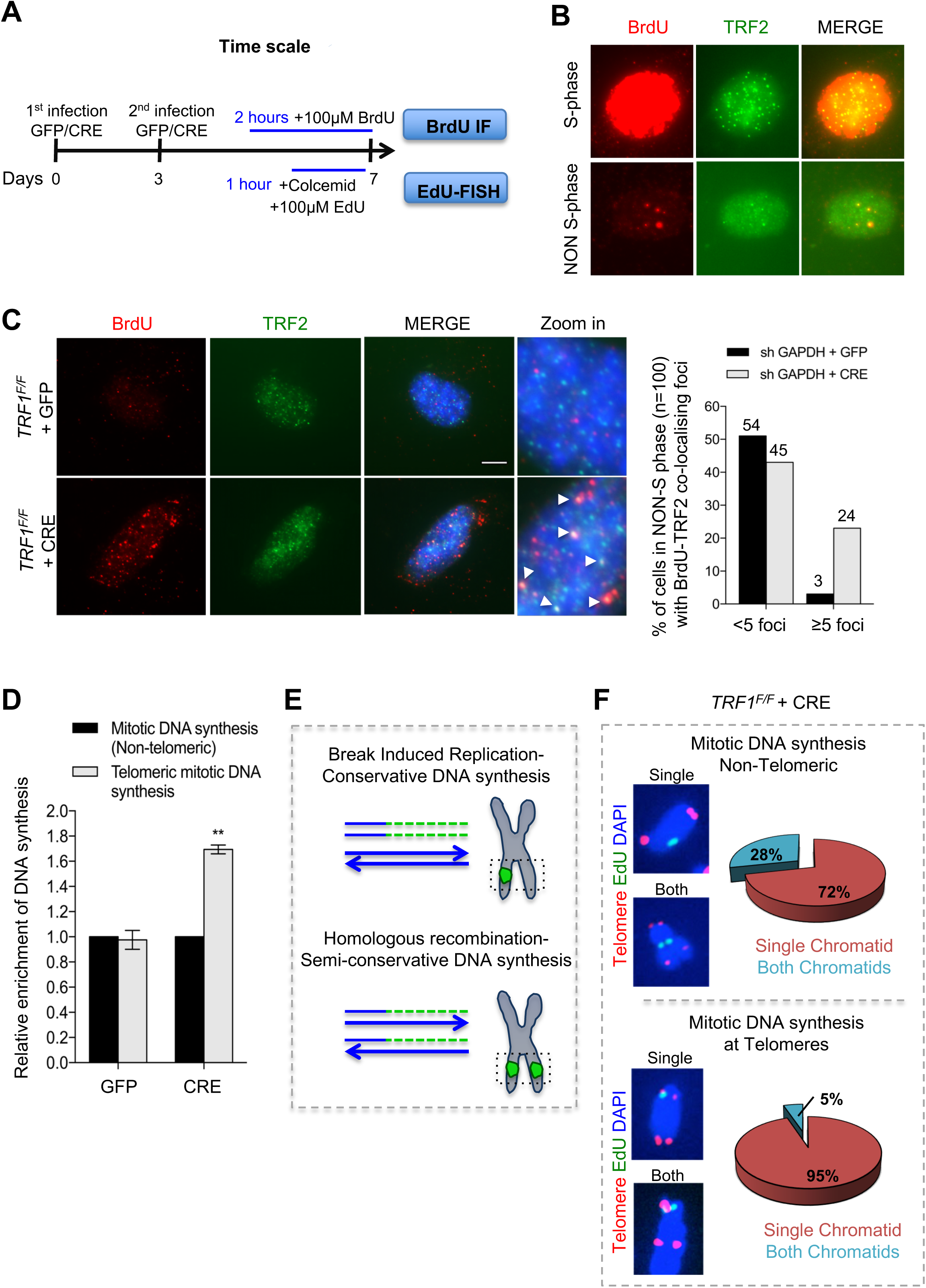
Deletion of *TRF1* induces mitotic DNA synthesis at telomeres. **(A)** Schematic overview of the experimental timeline. *TRF1^F/F^* MEFs cells were infected twice (day 0 and 3) with adenovirus containing either GFP control or CRE to mediate *TRF1* deletion. Prior to collection at day 7, cells were treated with either BrdU (100µM) for 2 hours or EdU (100µM) + colcemid for 1 hour, to perform respectively BrdU-Immunofluorescence (IF) or EdU-FISH on metaphases. **(B)** Representative image of BrdU (red) -TRF2 (green) immunofluorescence showing example of cells in S-phase (upper panel) and non-S-phase (bottom panel). (**C)** Immunofluorescence showing co-localisation of BrdU (red) with TRF2 (green) in *TRF1^F/F^* MEFs nuclei (DAPI, blue) treated with GFP and CRE. Scale bar, 5 µm (denaturing conditions). Data are represented as % of cells in non-S-phase showing < 5 or ≥ 5 BrdU-TRF2 co-localising foci (n=100 nuclei). **(D)** Quantification of DNA synthesis using the number of EdU-positive intra-chromosomes or telomeres in *TRF1^F/F^* cells infected with GFP and CRE relative to the GFP control (n=50 metaphases). Data are represented as relative enrichment to the GFP control ± SEM. P values, two-tailed student t-test (**, *P* < 0.01). **(E)** Schematic representation of Break Induced Replication (top part) with single EdU foci at a single chromatid and Homologous recombination (bottom part) with EdU foci at both chromatids. **(F)** Analysis of DNA synthesis in *TRF1* deleted cells. *Upper panel:* Non-telomeric mitotic DNA synthesis. Representative images showing EdU signal (green) in a single chromatid or in both chromatids. Pie chart representing % of chromosomes having EdU signal at a single chromatid or at both chromatids. *Bottom panel:* Telomeric mitotic DNA synthesis. Representative images showing EdU signal (green) at telomeres (red) at single or both chromatids. Pie chart representing % of chromosomes having EdU signal at telomeres at a single chromatid or both chromatids. Source data are provided as a Source Data File.

### Mitotic DNA synthesis at replication stressed telomeres is POLD3 dependent

BIR is a recombination dependent process reinitiating DNA replication when one end of a chromosome shares homology with the template DNA, leading to conservative DNA synthesis, which is dependent on RAD52 and POLD3 (*pol32* homolog in yeast) (Bhowmick, Minocherhomji, & Hickson, 2016; Sotiriou et al., 2016). ALT telomeres have recently been reported to be elongated by BIR, in a POLD3 and SMC5-dependent manner (Dilley et al., 2016; Min et al., 2017; Potts, Porteus, & Yu, 2006). Since the SMC5/6 complex was exclusively enriched in PICh purified TRF1 depleted telomeres (Figure 2A), we further investigated the role of POLD3 and SMC5 in BIR DNA synthesis observed in *TRF1^-/-^* MEFs. We generated *TRF1 ^F/F^* cells deficient in *SMC5* or *POLD3* using specific shRNAs. Upon infection with GFP or CRE adenovirus, we produced respectively single or double deletion *TRF1-SMC5* or *TRF1-POLD3* cell lines. Loss of SMC5 and TRF1 expression were confirmed by immunoblotting (Figure 5A-B), while mRNA levels of POLD3 were analysed by RT-QPCR (Figure 5C). We first confirmed that these deletions did not elicit a cell cycle arrest. We only noticed a slight decrease in population doublings in the double mutants, while all cell lines were still able to properly divide and incorporate EdU (Figure S5A-B). Thus, we carried out EdU-FISH in these cells to check for the presence of BIR (Figure 5D). We found that the enrichment of DNA synthesis at telomeres in *TRF1* deleted cells was suppressed in the double mutant *TRF1-POLD3*, while the double mutant *TRF1-SMC5* revealed similar telomeric DNA synthesis when compared to the single *TRF1* mutant (Figure 5E). First, these results confirm that BIR is the molecular mechanism taking place at TRF1 depleted telomeres. Second, SMC5 appears to be dispensable for BIR dependent DNA synthesis at these replication-stressed chromosome ends.

**Figure 5.**
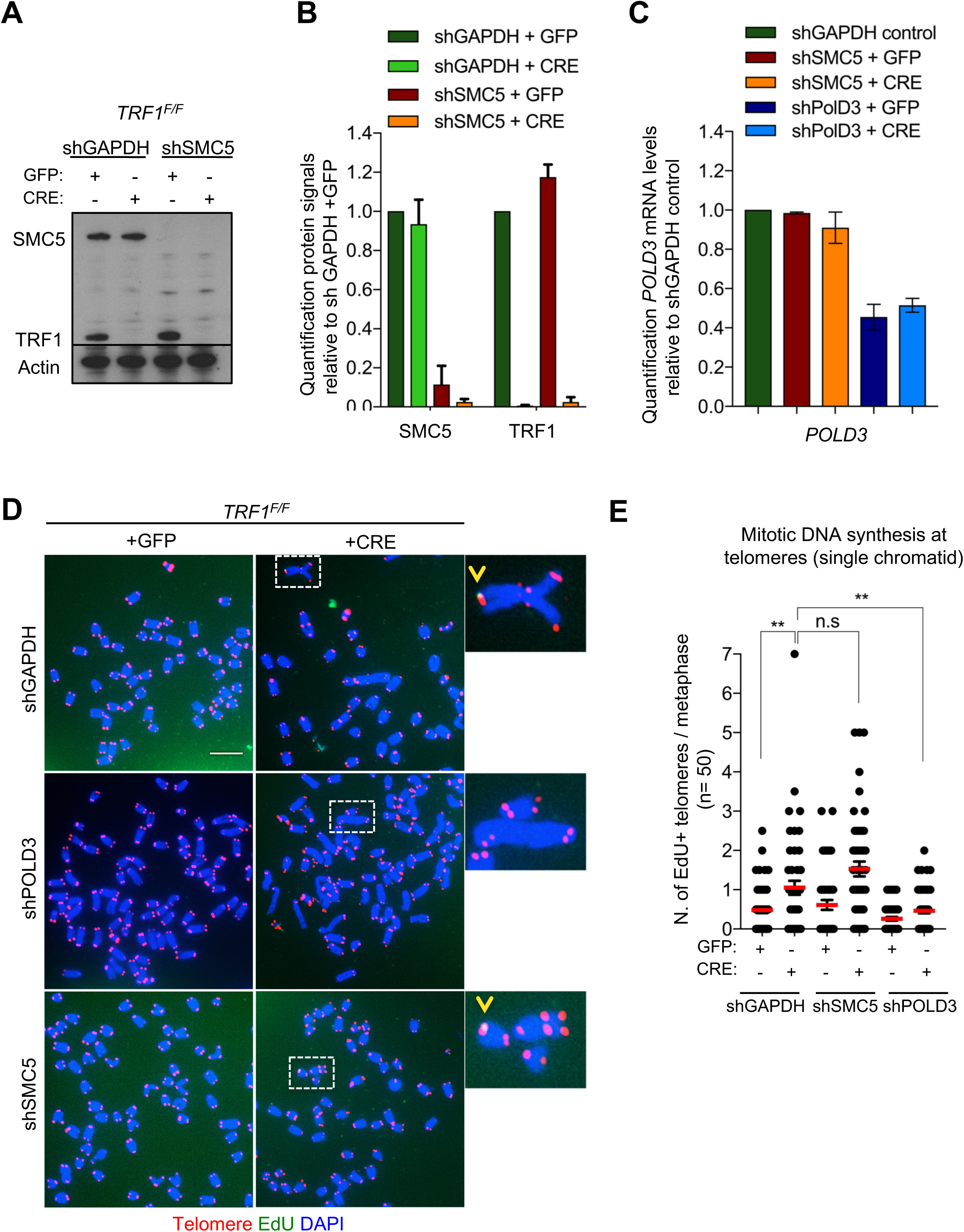
POLD3 but not SMC5 regulates mitotic DNA synthesis at *TRF1* deleted telomeres. **(A)** Western blotting showing expression of SMC5, TRF1 and Actin (loading control) proteins in *TRF1^F/F^* MEFs after infection with GFP or CRE-Adenovirus and deletion of *SMC5* by shRNA. shGAPDH is used as negative control. **(B)** Quantification of the knock-out and knock-down shown in A. Graph shows protein signal quantification relative to shGAPDH in +GFP control cells, data are represented as mean (n=3) ± SEM. **(C)** Quantification of *POLD3* mRNA levels relative to *GAPDH* control. Data are represented as mean (n=3) ± SEM. **(D)** Representative images of 6 different genotypes generated in the above description. Metaphases show EdU (green), telomeres labeled with TelPNA-C-rich-Cy3 (red) and chromosomes counterstained with DAPI (blue). Scale bar, 10 µm. **(E)** Quantification of mitotic DNA synthesis at telomeres (single chromatid) in *TRF1^F/F^* MEFs infected with shGAPDH control (GFP or CRE), shSMC5 (GFP or CRE) and shPOLD3 (GFP or CRE). Data are represented as number of EdU positive telomeres per metaphase ± SEM. n=50 metaphases. P value, two-tailed student t-test (**, *P* < 0.01; n.s.= non-significant). Source data are provided as a Source Data File.

### SMC5 and POLD3 are required for APBs formation and recombination at TRF1 deficient telomeres

We further examined whether POLD3 and SMC5 could be responsible not only for the BIR dependent DNA synthesis but also for the other ALT-like phenotypes observed at TRF1 deficient telomeres. Since TRF1 is well known to suppress telomere fragility or MTS (Sfeir et al., 2009) (Martinez et al., 2009), we first investigated the role of POLD3 and SMC5 in the induction or maintenance of this telomere replication stress in the double mutants (Figure S6A-B). As previously reported, TRF1 depleted telomeres present approximately 20% of fragile telomeres per chromosomes (Figure S6C). We could not detect any changes in the frequency of telomere fragility in *TRF1-POLD3* nor *TRF1-SMC5* mutants (Figure S6C) suggesting that neither POLD3 nor SMC5 are involved in the mechanism that gives rise to telomere fragility. As APBs were increased in *TRF1* deleted cells (Figure 2C), we investigated the roles of *POLD3* and *SMC5* in the formation of these specialised bodies. A significant reduction in number of cells having co-localising PML-telomere foci was detected in the double mutant cells *TRF1-POLD3* and *TRF1-SMC5* (Figure 6A) suggesting that *POLD3* and *SMC5* are necessary for the formation of these recombination machinery loci. We next explored the involvement of these two factors in HR by scoring for T-SCE (Figure 6B), discriminating also between the two categories of T-SCEs (single or double exchanges) in the analysis of the double mutants *TRF1-SMC5* and *TRF1-POLD3*. We found that both types of exchanges are dependent on SMC5 and POLD3 (Figure 6B-S6D). Finally, we assessed TERRA expression levels in the double mutants. Surprisingly, only the absence of *POLD3* was able to rescue the increase in TERRA levels detected in *TRF1* deficient cells, while the *SMC5* single mutant increased TERRA expression (Figure 6C). Collectively, our data indicate that both POLD3 and SMC5 are essential for T-SCE and APBs formation, but only POLD3 is required to maintain increased TERRA levels and BIR observed in *TRF1* deficient cells. This suggests that POLD3 and SMC5 have separate roles or act at different stages of the recombination events happening at TRF1 depleted telomeres, advocating also an intriguing connection between TERRA and BIR. We speculate that TERRA could trigger the homology search by stimulating the initial steps of BIR in which POLD3 is involved (Figure 7).

**Figure 6.**
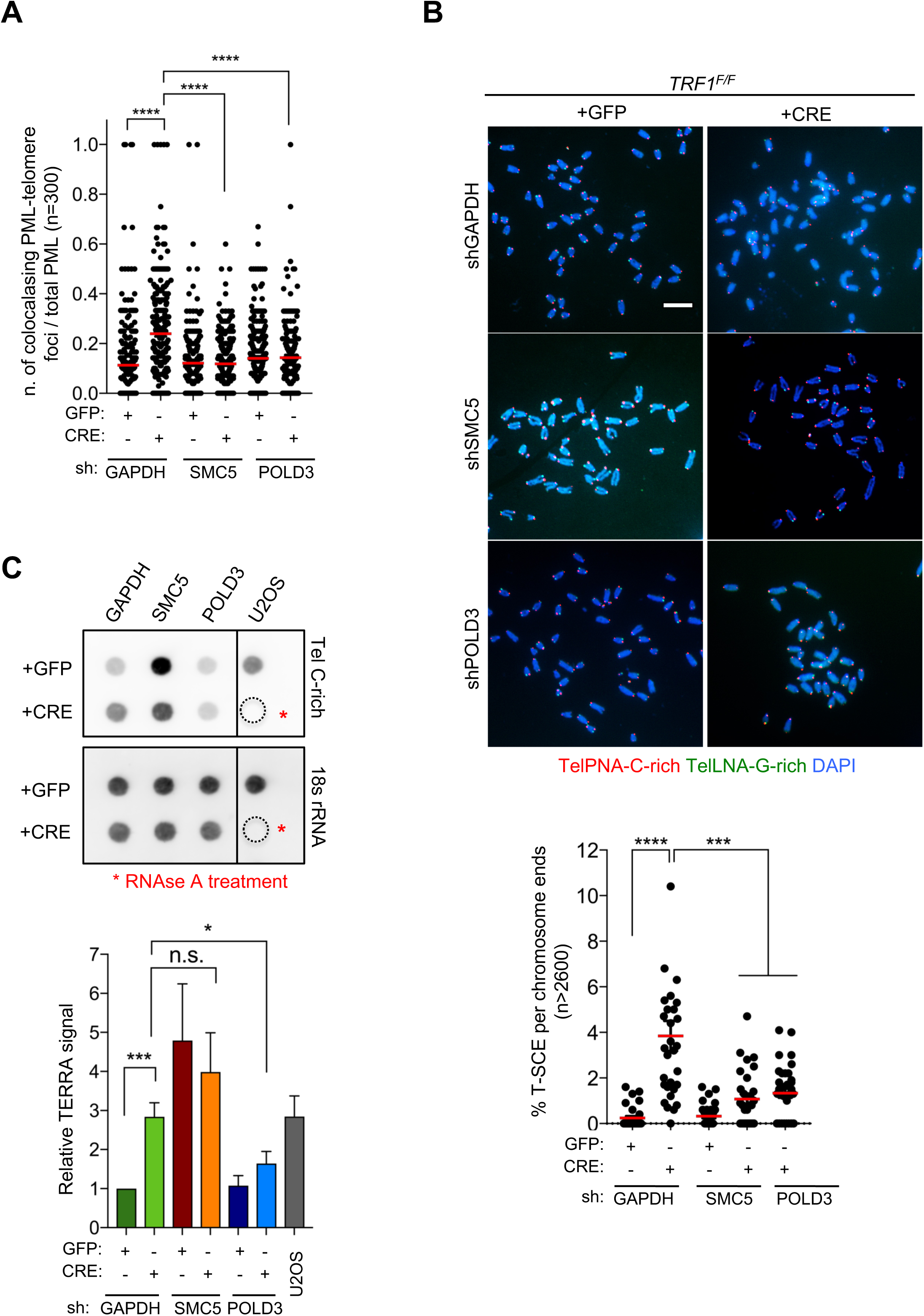
*SMC5* and *POLD3* are required for induction of recombination at *TRF1* deficient telomeres. **(A)** APBs formation in *TRF1* deleted cells is rescued in double mutants *TRF1-SMC5* and *TRF1-POLD3*. Quantification of APBs formation is represented as number of co-localising PML-telomere foci divided by the total number of PML present per nucleus (n=300 nuclei analysed) ± SEM. **(B)** Representative images of the chromosome oriented (CO)-FISH assay <u>with denaturation</u>, used to score for telomeric T-SCEs in *TRF1^F/F^* MEFs infected with shGAPHH control (GFP or CRE), shSMC5 (GFP or CRE) and shPOLD3 (GFP or CRE). Telomeres are labeled with TelPNA-C-rich-Cy3 (red) and TelLNA-G-rich-FAM (green), while chromosomes are counterstained with DAPI (blue). Scale bar, 10 µm. For quantification T-SCE was considered positive when involved in a reciprocal exchange of telomere signal with its sister chromatid (both telomeres yellow) and for asymmetrical exchanges at single chromatid (one telomere yellow). Data are indicated as % of T-SCE per sister telomere (bottom panel). The mean values (n=>2600 chromsome ends) ± SEM are indicated. P value, two-tailed student t-test (***, *P* < 0.001; ****, *P* < 0.0001). **(C)** RNA dot blot analysis in *TRF1*, *SMC5, POLD3* single and double mutants. The blot was revealed with a DIG-Tel-C-rich probe or 18s rRNA as a control. TERRA signals were normalised to 18s rRNA and GFP control (bottom panel). Data are represented as relative TERRA signal (n = 4) ± SEM. P values, two-tailed student t-test (*, *P* < 0.05; ***, *P* < 0.001; n.s.= non-significant). Source data are provided as a Source Data File.

**Figure 7.**
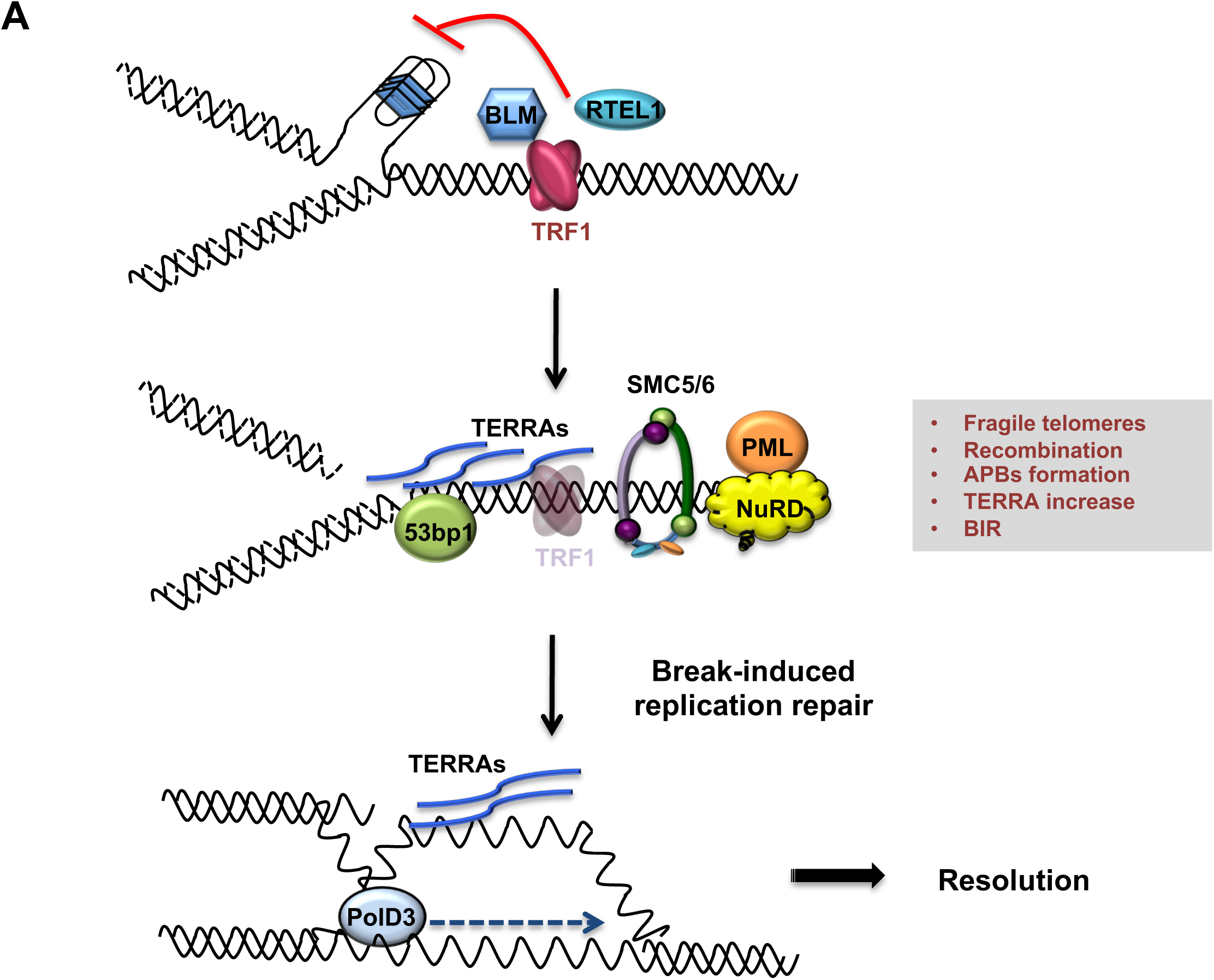
Model describing TRF1 as a negative regulator of telomeric transcription (TERRAs), APBs formation, telomeric recombination via PolD3-BIR dependent pathway. Replicative stress induced by TRF1 deletion alters the chromatin status of these telomeres. Recruitment of chromatin remodelers/HR factors, TERRA accumulation and telomere fragility are observed. The SMC5/6 complex and polymerase POLD3 are among the factors recruited at replicative-stressed telomeres, representing the key players for APBs formation and telomere recombination, particularly BIR-mechanism. We propose that increased TERRAs molecules at telomeres could lead to increased R-loops, which are bypassed by POLD3 dependent BIR to resolve fork progression hindrance.

## Discussion

Faithful DNA replication of genetic information is essential for the maintenance of genome stability and integrity. Specific genomic loci, including fragile sites and telomeres, represent major obstacles to DNA replication progression and/or completion. Fragile sites have the propensity to form visible gaps or breaks on chromosome in metaphase spreads of cell lines from patients having fragile X-syndrome or Huntington’s disease (reviewed in(Minocherhomji & Hickson, 2014; Minocherhomji et al., 2015)). It is well documented that CFS expression is exacerbated in cells grown under low to mild replication stress, for example upon inhibition of DNA polymerase with APH (Minocherhomji & Hickson, 2014; Minocherhomji et al., 2015). Fragile sites are hotspots for deletions, chromosome rearrangements and are associated with an increased frequency of homologous recombination (Glover & Stein, 1987). Over the last decade, telomeres have been identified as APH induced fragile sites displaying the standard phenotype of multiple spatially distinct telomere foci (MTS or telomere fragility) on metaphase spreads (Martinez et al., 2009; Sfeir et al., 2009). Various factors suppress MTS and thereby facilitate DNA replication at telomeres; including (Zaaijer, Shaikh, Nageshan, & Cooper, 2016) the shelterin protein TRF1 (Martinez et al., 2009; Sfeir et al., 2009), the DNA helicases RTEL1 (Vannier et al., 2012), BLM and WRN (Barefield & Karlseder, 2012), Topoisomerase TopoIIa (d’Alcontres, Palacios, Mejias, & Blasco, 2014) and Rif1 (Zaaijer et al., 2016). High levels of DNA damage and telomere fragility are characteristics of ALT cells (Cesare et al., 2009) (Min et al., 2017), presenting several DNA repair and damage factors in APBs (Draskovic et al., 2009; G. Wu et al., 2000; Yeager et al., 1999), indication of elevated telomeric stress in these cells. Therefore, it has been hypothesized that ALT mechanism arises from persistent replication stress, which can be resolved by the initial collapse of the replication fork, subsequently offering substrates for HR repair mechanisms dependent on homology search and telomere synthesis as reported with BIR pathway (Dilley et al., 2016).

In this study, we report that replication stress generated at TRF1 depleted telomeres in telomerase positive MEFs is associated with the recruitment of ALT signature factors including PML, subunits of the NuRD complex, BRCA1 and SMC5/6 complex. We suggest that the formation of permissive telomeric chromatin enables transcription of telomeric sequences into TERRAs and increases recombination as measured by T-SCEs, in a POLD3 and SMC5/POLD3 dependent manner, respectively. Moreover, we detect mitotic DNA synthesis at TRF1 depleted telomeres, which is dependent on POLD3 but not SMC5. Collectively, the presence of replication stress, recombination, APBs formation, TERRA increase and recruitment of specific chromatin factors, suggest a strong analogy between MEFs telomeres deleted for TRF1 and ALT telomeres, supporting the hypothesis that replicative stress could be the source of ALT initiation.

We suggest that chromatin remodeling factors such as NuRD-ZNF827 are recruited to TRF1 deficient telomeres to counteract the shelterin instability. This may be explained by analogy with ALT telomeres where telomeric DNA sequence is interspersed with variant repeats (Conomos et al., 2012; Marzec et al., 2015), which are suggested to cause displacement of shelterin proteins (Conomos, Pickett, & Reddel, 2013), thus increasing replication stress and DDR. In this scenario, nuclear receptors bind the interspersed variant repeats and recruit several chromatin remodeling factors including the NuRD complex, which can further alter the telomere architecture by increasing telomere compaction (Conomos et al., 2014); perhaps a transient state before stimulating telomere associations and generating more ‘open’ recombination permissive conditions at telomeres. In line with our findings, repressive chromatin at DSBs has been proposed to facilitate homology search and promote recruitment of HR proteins like BRCA1 (Khurana et al., 2014). In addition to BRCA1, we have identified through PICh analysis the SMC5/6 complex specifically recruited at *TRF1* deficient telomeres. We demonstrate that this complex plays the same role at replication induced telomeres as in ALT cells, targeting telomeres to PML bodies (APBs) and facilitating telomeric HR at these sites (Potts et al., 2006), since double mutant *SMC5-TRF1* disrupts formation of APBs and reduces T-SCEs events. However, we were unable to fully induce ALT in *TRF1* deficient MEFs, as they display neither C-circles nor heterogeneity in telomere length and telomerase is still active. The latter could act as a stabiliser of telomeric DNA ends generated during fork restart (Tong et al., 2015), similarly to what happens in *RTEL1^-/-^* MEFs (Margalef et al., 2018) and in human *RTEL1* deficient cells with long telomeres (Porreca et al., 2018).

Persistent DNA damage in ALT cells is suggested to originate from telomeric replication stress, which is proposed to be resolved by BIR in a POLD3 dependent manner (Dilley et al., 2016; Min et al., 2017; Roumelioti et al., 2016). Our results show that TRF1 is a major suppressor of telomeric replication stress and consequently of POLD3 dependent BIR. *TRF1* deficient telomeres present slower movement of S-phase replication forks, measured by molecular combing (Sfeir et al., 2009). The slower replication rates at telomeres is proposed to be a consequence of the hindrance of the replication forks by DNA secondary structures, including formation of G-quadruplexes on the lagging strand template or RNA-DNA hybrids. In the absence of TRF1, BLM is unable to be recruited to replicated telomeres and to open DNA secondary structures (Lee et al., 2018; Zimmermann, Kibe, Kabir, & de Lange, 2014). Based on our results, we propose that in the absence of TRF1, POLD3 dependent BIR bypasses the stalled replication fork during G2/M phase.

Recent studies identified BIR as a mechanism to bypass RNA-DNA hybrids in a Rad52 and Pol32 dependent manner in yeast (Amon & Koshland, 2016; Neil, Liang, Khristich, Shah, & Mirkin, 2018). This mechanism is also conserved in human cells where POLD3 is necessary for the restart of stalled replication forks at RNA-DNA hybrids (Tumini, Barroso, Calero, & Aguilera, 2016). Altogether, we propose that increased TERRAs levels at TRF1 depleted telomeres could form RNA-DNA hybrids that are bypassed by POLD3 dependent BIR (Figure 7). This is in agreement with recent findings showing that TRF1 suppresses R-loop formation mediated by TRF2 (Lee et al., 2018). In contrast to Pold3, SMC5 acts as inhibitor of TERRA accumulation, as its absence is causing a significant increase in TERRA levels. This result is reminiscent of the role of yeast Smc5 in facilitating the resolution of toxic recombination intermediates at RNA-DNA hybrids generated by the helicase Mph1 (Chen et al., 2009; Lafuente-Barquero et al., 2017). We also describe a role of SMC5 in promoting T-SCEs, but not MiDAS formation (in contrast to PolD3), in the absence of TRF1. These results are indicating an exclusive function of SMC5 in HR at replicative-stressed telomeres, perhaps ensuring the right balance between accumulation and removal of HR-dependent intermediates formed during DNA repair (Aragon, 2018). On the other hand, the lack of Smc5 in promoting MiDAS seems in apparent contradiction with a recent observation in ALT cells (Min et al., 2017), where telomeric MiDAS is decreased in SMC5/6-depleted Saos2 cells. We speculate, this difference is due to an imbalance of factors used for ALT maintenance, compared to the early events observed in our conditional system after only few population doublings. Therefore, we cannot rule out a possible role of SMC5/6 in promoting MiDAS at a later stage, similar to the one observed in ALT maintenance.

Along with MUS81 structure specific nuclease, POLD3 and POLD4 subunits of the DNA polymerase delta are essential for CFS expression observed in human cells under replication stress (Minocherhomji et al., 2015; Tumini et al., 2016). To our surprise, *TRF1-POLD3* double mutant did not show any suppression of telomere fragility, indicating key differences in the mechanism generating these phenotypes.

In conclusion, our analysis of TRF1 function provides a molecular understanding of the level of protection that this shelterin protein offers at telomeres. The role of TRF1 in facilitating DNA replication at telomeres was already described but only until a certain extent. Surprisingly, we establish that TRF1 is essential for the suppression of early ALT-like signature events including heterochromatin remodeling, telomeric transcription (TERRAs), APBs formation and increased POLD3-BIR dependent telomeric recombination.

## Online Methods

### Cell culture, viral transductions and transfections with siRNAs

TRF1 conditional knock-out MEFs (SV40-immortalised) were described previously (Martinez et al., 2009; Sfeir et al., 2009). Cells were cultured at 37°C in 5%CO2, using DMEM medium supplemented with 10% FCS (Sigma F2442). To achieve TRF1 deletion, cells were infected twice at 72h interval with Ad5-CMV-CRE (m.o.i. of 50) and harvested 3 or 4 days after the second infection.

pLKO.1-puromycin lentiviral vectors containing shRNAs for SMC5 (sequence CCCATAATGCTCACGATTAAT, Sigma), POLD3 (GCATATACTCATGTGTGGTTT, Dharmacon) or GAPDH (CTCATTTCCTGGTATGACA, Open biosystems) were introduced by infection of lentivirus-containing supernatant from 293FT cells. Puromycin selection was performed for 3 weeks at 2µg/ml and several clones were expanded and cultured before screening them for knock-down efficiency.

### Western blot

Cells were scraped in cold PBS, spun down and incubated in lysis buffer (NaCl 40 mM; Tris 25 mM, pH 8; MgCl 2 mM; SDS 0.05%; Benzonase 1µl/2ml; Complete protease inhibitor cocktail, EDTA-free, Roche) for 10 min on ice. The lysates were sheared 10 times by forcing it through a 25G needle and left on ice for another 10 min. 35 μg of protein lysates were denatured for 10 min at 95°C after addition of Laemmli buffer 4X (50mM Tris pH7; 100mM DTT; 2%SDS; 0.1%bromophenol blue; 10% glycerol), separated on 4-12% Bis-Tris gels (Invitrogen) and transferred onto a nitrocellulose membrane (Amersham Protran 0.2µm NC). Rabbit anti-TRF1 (gift from Titia de Lange) and rabbit anti-SMC5 (gift from Jo Murray) antibodies were diluted in PBST (PBS1x; 0.1 % Tween-20, Sigma-Aldrich) with 5% non-fat milk. Following incubations with HRP-coupled secondary antibodies signals were visualised using ECL II kit (Pierce) and x-ray film exposure (Amersham Hyperfilm ECL). Beta-actin antibody was used for normalisation (Abcam, ab8226).

### Quantitative RT-PCR

RNA extraction was carried out using RNeasy Mini Kit (Qiagen). 500ng of RNA were subjected to reverse transcription using random hexamer primers and cDNA Synthesis Kit (Roche) according to the manufacturer’s protocol. Quantitative PCR was performed using QuantiTect SYBR Green PCR Master Mix and the following primers: mouse POLD3 with antisense 5’-ACACCAAGTAGGTAACATGCAG-3’ and sense 5’-AAGATCGTGACTTACAAGTGGC-3’ sequences; Mouse Actin with antisense 5’-CCAGTTGGTAACAATGCCATGT-3’ and sense 5’-GGCTGTATTCCCCTCCATCG-3’ sequences; The PCR cycles were as follows: 95°C for 15 min, 95°C for 15 sec, 55°C for 30 sec, 72°C for 30 sec for 44 cycles.

### Telomeric Chromatin Isolation by PICh

PICh was carried out as previously described (Dejardin & Kingston, 2009) using the following 2’Fluoro-RNA probes for hybridisation: Destiobiotin-108 atom tether-UUAGGGUUAGGGUUAGGGUUAGGGt (Telo probe); Destiobiotin-108 atom tether-GAUGUGGAUGUGGAUGUGGAUGUGg (Scramble probe).

### Gel & post digestion processing

Gels were processed using a variant of the in-gel digestion procedure as described in (Shevchenko, Tomas, Havlis, Olsen, & Mann, 2006). Briefly, gel sections were excised and chopped into uniform cubes, followed by de-staining with 50/50, 50mM ammonium bicarbonate (AmBic)/acetonitrile(ACN). Gel sections were then dehydrated with 100% ACN followed by the subsequent sequential steps: reduction with 10mM dithiothreitol (DTT) at 56°C for 30 minutes in the dark, dehydration, alkylation with 55mM iodoacetamide (IAM) at RT for 20 minutes in the dark and dehydration. Gel sections were finally re-hydrated with a 40mM AmBic, 10% ACN solution containing 500ng of Trypsin Gold (Promega, V5280) and incubated overnight at 37°C. Recovered gel digest extracts were dried on a speed-vac, reconstituted with 99/1, H2O/ACN + 0.1% FA and de-salted using a standard stage tip procedure using C18 spin tips (Glygen Corp, TT2C18). Dried gel digest peptide extracts solubilised in 25µl of 0.1% trifluoroacetic acid (TFA) and clarified solution transferred to auto sampler vials for LC-MS analysis.

### Mass spectrometry analysis

Peptides were separated using an Ultimate 3000 RSLC nano liquid chromatography system (Thermo Scientific) coupled to a LTQ Velos Orbitrap mass spectrometer (Thermo Scientific) via an EASY-Spray source. 6μL of sample was loaded in technical duplicates onto a trap column (Acclaim PepMap 100 C18, 100μm × 2cm) at 8μL/min in 2% acetonitrile, 0.1% TFA. Peptides were then eluted on-line to an analytical column (EASY-Spray PepMap C18, 75μm × 25cm). Peptides were separated using a linear 120 minute gradient, 4-45% of buffer B (composition of buffer B– 80% acetonitrile, 0.1% formic acid). Eluted peptides were analysed by the LTQ Velos operating in positive polarity using a data-dependent acquisition mode. Ions for fragmentation were determined from an initial MS1 survey scan at 15000 resolution (at m/z 200), followed by Ion Trap CID (collisional induced dissociation) of the top 10 most abundant ions. MS1 and MS2 scan AGC targets set to 1e6 and 1e4 for a maximum injection time of 500ms and 100ms respectively. A survey scan m/z range of 350 – 1500 was used, with a normalised collision energy set to 35%, charge state rejection enabled for +1 ions and a minimum threshold for triggering fragmentation of 500 counts.

### Data analysis

All data files acquired were loaded into MaxQuant(J. Cox et al., 2014) version 1.6.0.13 analysis software. Raw files were combined into an appropriate experimental design to reflect technical and biological replicates. The LFQ algorithm and match between runs settings were selected. Data were searched against the UniProt Reference Proteome Mus musculus protein database (UP000000589), downloaded on 16th January 2019 from the UniProt website. The database contains 17,002 reviewed (Swiss-Prot) & 37,186 un-reviewed (TrEMBL) protein sequences. MaxQuant also searched the same database with reversed sequences so as to enable a 1 % false discovery rate at peptide and protein levels. A built-in database of common protein contaminants was also searched.

Upon completion of the search, the “proteingroups.txt” output file was loaded in Perseus version 1.4.0.2. Contaminant and reverse protein hits were removed. LFQ intensities were log2 transformed. Data were group categorised to “Scramble”, “Telomere” or “Deletion”. Data were filtered for a minimum of 3 valid LFQ intensity values in at least one group. Missing values (NaN) were imputed from a normal distribution with default values.

### FISH and CO-FISH on metaphase spreads

For metaphase spread preparation, cells were incubated for 60 minutes with 10ng/ml colcemid (Roche). Cells were harvested, swollen in 75 mM KCl solution for 15 min at 37°C, fixed in ethanol/acetic acid solution (3:1, v/v) and washed three times with the same fixing solution. Suspensions of fixed cells were dropped onto glass slides and dried overnight before performing FISH experiments.

Q-FISH and CO-FISH procedures were performed as previously described (Ourliac-Garnier & Londono-Vallejo, 2011). Briefly, metaphase spreads were fixed in 4% formaldehyde for 2 min, washed 3 × 5 min in PBS 1x, treated with pepsin (1 mg/ml in 0.05 M citric acid pH 2) for 10 min at 37°C, post-fixed for 2 min, washed and incubated with ethanol series (70%, 80%, 90%, 100%). Hybridising solution containing Cy3-O-O-(CCCTAA)3 probe (PNA bio) in 70% formamide, 10 mM Tris pH 7.4 and 1% blocking reagent (Roche, 11096176001) was applied to each slide, followed by denaturation for 3 min at 80°C on heating block. After 2 hour hybridisation at RT, slides were washed twice 15 min in 70% formamide, 20 mM Tris pH 7.4, followed by three washes of 5min in 50 mM Tris pH 7.4, 150 mM NaCl, 0.05% Tween-20, dehydrated in successive ethanol baths and air-dried. Slides were mounted in antifade reagent (ProLong Gold, Invitrogen) containing DAPI and images were captured with Zeiss microscope using Carl Zeiss software. Telomeric signals were quantified using the ImageJ software (Fiji).

For CO-FISH, the cells were treated with 10µM BrdU:BrdC (3:1) for 16h, followed by colcemid treatment as above. Prior to hybridisation slides were treated with RNAse A (0.5µg/ml in PBS) for 10 min at 37°C, incubated with Hoechst (1 µg/ml in 2XSSC) for 10 min at RT, exposed to UV light for 1h and treated with *Exo*III to degrade the neosynthesised DNA strand containing BrdU/C. Slides were next dehydrated through ethanol series, hybridising solution containing TelG-FAM probe (Exiqon) in 50% formamide, 2XSSC, 1% blocking reagent was applied to each slide, followed by denaturation for 3 min at 80°C on heating block and hybridisation for 2 hours in the dark. Slides were washed 2 x 15 min in 50% formamide, 2XSSC and 3 × 5 min in 50 mM Tris pH 7.4, 150 mM NaCl, 0.05% Tween-20. Finally, slides were dehydrated, incubated with TelC-cy3 probe for 2 hours, followed by the steps described above in the FISH protocol.

### Immunofluorescence-FISH

Cells seeded on slides were permeabilised with Triton X-100 buffer (0.5% Triton X-100; 20mM Tris pH8; 50mM NaCl; 3mM MgCl2; 300mM sucrose) at RT for 5min and then fixed in 3% formaldehyde/2%sucrose in PBS1X for 15min at RT and washed three times in PBS1X. After a 10 min permeabilisation step and a wash in PBS1X, nuclei were incubated with blocking solution (10% serum in PBS1X) for 30 min at 37°C and stained with specific primary antibodies: rabbit anti-PML (1/200, a gift from Paul Freemont); rabbit anti-53bp1 dilution (1/400, Bethyl A300-272A). After three washes in PBS1X, nuclei were incubated with secondary donkey anti-rabbit Alexa 488 antibody (1/400, Life Technologies) for 40 min at 37°C, washed three times in PBS1X, post fixed 10 min and hybridised with TelC-cy3 PNA probe as described in FISH protocol.

EdU labeling and staining were performed as previously reported (Minocherhomji et al., 2015). Briefly, cells were incubated 1h with EdU (100µM) and colcemid (10ng/ml), followed by metaphase spread preparation. For EdU staining, the steps of fixation, pepsin treatment and dehydration in ethanol serial dilutions were carried out as in FISH protocol, followed by Click IT assay using EdU-Alexa Fluor 488 imaging kit according to the manufacturer’s instructions (Thermo Fisher). Metaphases were post-fixed and hybridised with TelC-cy3 PNA probe.

### TERRA-FISH

TERRA-FISH experiment was carried out as previously described (Azzalin, Reichenbach, Khoriauli, Giulotto, & Lingner, 2007) with minor modifications. Briefly cells were permeabilised 5 min with cold CSK buffer (10mM Pipes pH7;100mM NaCl; 300 mM sucrose; 3mM MgCl2; 0.5% Triton X-100 and 10mM of inhibitor Ribonucleoside Vanadyl Complex). After a wash in PBS1X, cells were fixed for 10 min in 3%formaldeyde solution and washed three times with PBS, followed by Immunofluorescence with primary anti-TRF2 (dilution 1/10.000, 1254 ab gift from T. de Lange). Nuclei were then incubated with secondary donkey anti-rabbit Alexa 488 antibody (1/400, Life Technologies) for 40 min at 37°C, washed three times in PBS1X and post fixed for 10 min. After incubation with ethanol series (70%, 80%, 90%, 100%) slides were dried O/N in the dark. TelC-cy3 PNA probe was used for TERRA detection and after incubation for 2hours at RT, slides were washed 3 x 5 min in 50% formamide, 2XSSC at 39°C, 3 × 5 min in 2XSSC at 39°C and a final wash in 2XSSC at RT. Slides were dehydrated in successive ethanol baths, air-dried and mounted in antifade reagent (ProLong Gold, Invitrogen) containing DAPI and images were captured with Zeiss microscope using Carl Zeiss software. Quantification was performed using CellProfiler 3.1.8 software.

### Chromatin Immunoprecipitation (ChIP)

Chromatin preparation and ChIP experiments were performed as previously described (Porreca et al., 2018) with the following modifications: sonication of chromatin was performed for 20 min (30 sec on / 30 sec off) in a Diagenode water bath-sonicator at high speed. 20-50 μg of chromatin was diluted 10 times in ChIP dilution buffer (20 mM Tris-HCl pH 8, 150 mM KCl, 2 mM EDTA pH 8, 1% Triton X-100, 0.1% SDS), pre-cleared with Dynabeads (Invitrogen) and incubated overnight with 2-5 μg of antibody (listed in Table S1).

### RNA dot blot

RNA extraction was carried out using RNeasy Mini Kit (Qiagen), according to the manufacturer instructions. 2µg of RNA were denatured in 0.2 M NaOH by heating at 65°C for 10 min, incubated 5min on ice and spotted onto a positively charged Biodyne B nylon membrane (Amersham Hybond, GE Healthcare). Membranes were UV-crosslinked (Stratalinker, 2000 kJ) and baked for 45 min at 80°C, followed by hybridisation at 42°C with digoxigenin (DIG)-labeled telomeric C-rich oligonucleotide TAA(CCCTAA)4, prepared using 3’ end labeled kit (Roche). Signal was revealed using the anti-DIG-alkaline phosphatase antibodies (Roche) and CDP-Star (Roche) following the manufacturer’s instructions. Images were captured using the Amersham Imager 680 (GE Healthcare) and analysed using the Image Studio Lite software. 18s rRNA probe with sequence: 5’-CCATCCAATCGGTAGTAGCG was used for normalisation.

### Northern Blot

10µg of RNA was denatured for 10 min at 65°C in sample buffer (50% formamide, 2.2M formaldehyde, 1X MOPS) followed by ice incubation for 5 min. 10X Dye buffer (50% Glycerol, 0.3% Bromophenol Blue, 4mg/ml Ethidium Bromide) was added to each sample and all of them were run on a formaldehyde agarose gel (0.8% agarose, 1X MOPS, 6.5% formaldehyde) at 5V per cm in 1X MOPS buffer (0.2M MOPS, 50mM NaOAc, 10 mM EDTA, RNAse free water). The gel was rinsed twice in water, washed twice with denaturation solution (1.5M NaCL, 0.05M NaOH), followed by additional three washes with 20XSSC before transferring the RNA on a positively charged Biodyne B nylon membrane (Amersham Hybond, GE Healthcare) using a neutral transfer in 20XSSC. The membrane was fixed and detected as described for the RNA dot blot.

## Acknowledgements

We thank Titia de Lange for providing TRF1 antibody, Paul Freemont for providing PML antibody and Jo Murray for SMC5 antibody. Special thanks to Paulina Marzec for teaching R.M.P the PICh techniques in Boulton Lab. We thank Arturo Londono, Julia P. Cooper, Julian Sale, Titia de Lange, Valerie Borel, Simon Boulton for their helpful comments. TRF1 MEFs were generated in Simon Boulton’s group from Titia de Lange’s mouse line and Clare H. McGowan’s group, respectively. Vannier lab work is supported by the London Institute of Medical Sciences (LMS), which receives its core funding from UKRI (previously MRC) and by an ERC Starter Grant (637798; MetDNASecStr).

## Author contributions

R.M.P and J.B.V designed the project and wrote the manuscript. R.M.P, P.P.L, E.H.M, R.G.F and J.B.V conducted experiments. A.M, P.F, H.K performed mass spectrometry analysis.

## Data availability statement

All relevant data are available from the authors. The source data underlying Figs 1B, 1D, 1F, 2B-2D, S2, 3A-C, 4C, 4D, 4F, 5A-C, 5E, 6A-C and supplementary Figs S1A-S1B, S3B-S3D, S4A-S4C, S5A,-S5C and S6D are provided as a Source data file. Accession code for the proteomic data will be made available before publication on public repository PRIDE.

---

**Table S1.**
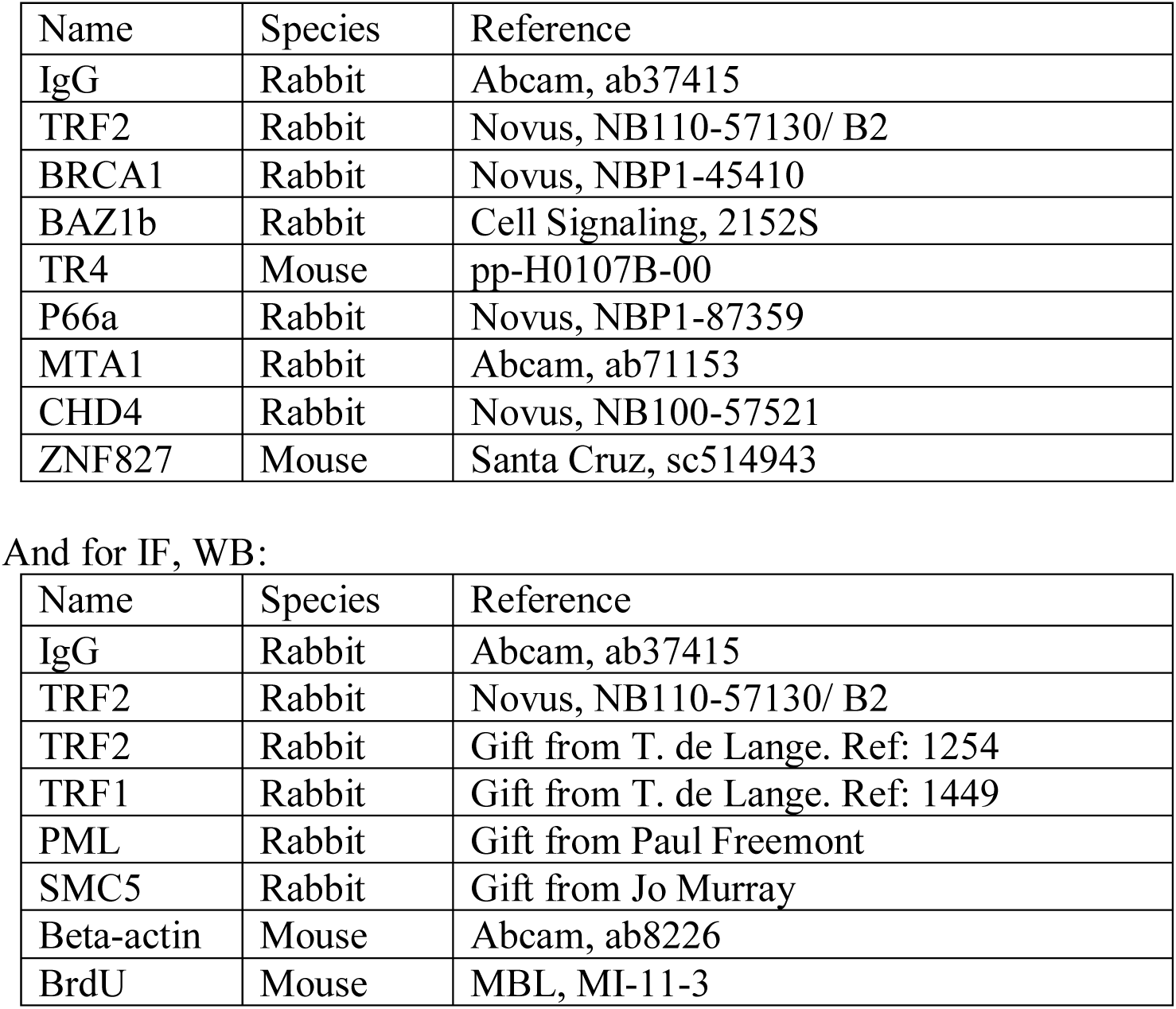
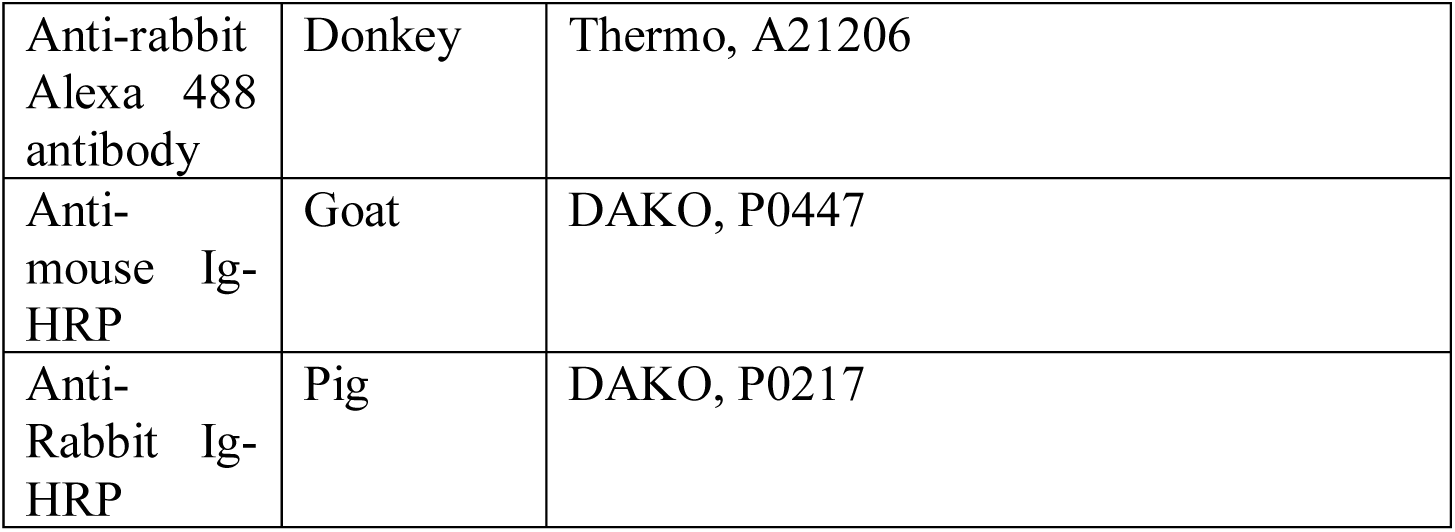
List of antibodies used for ChIP

## Supplemental Figure legends

**Figure S1.**
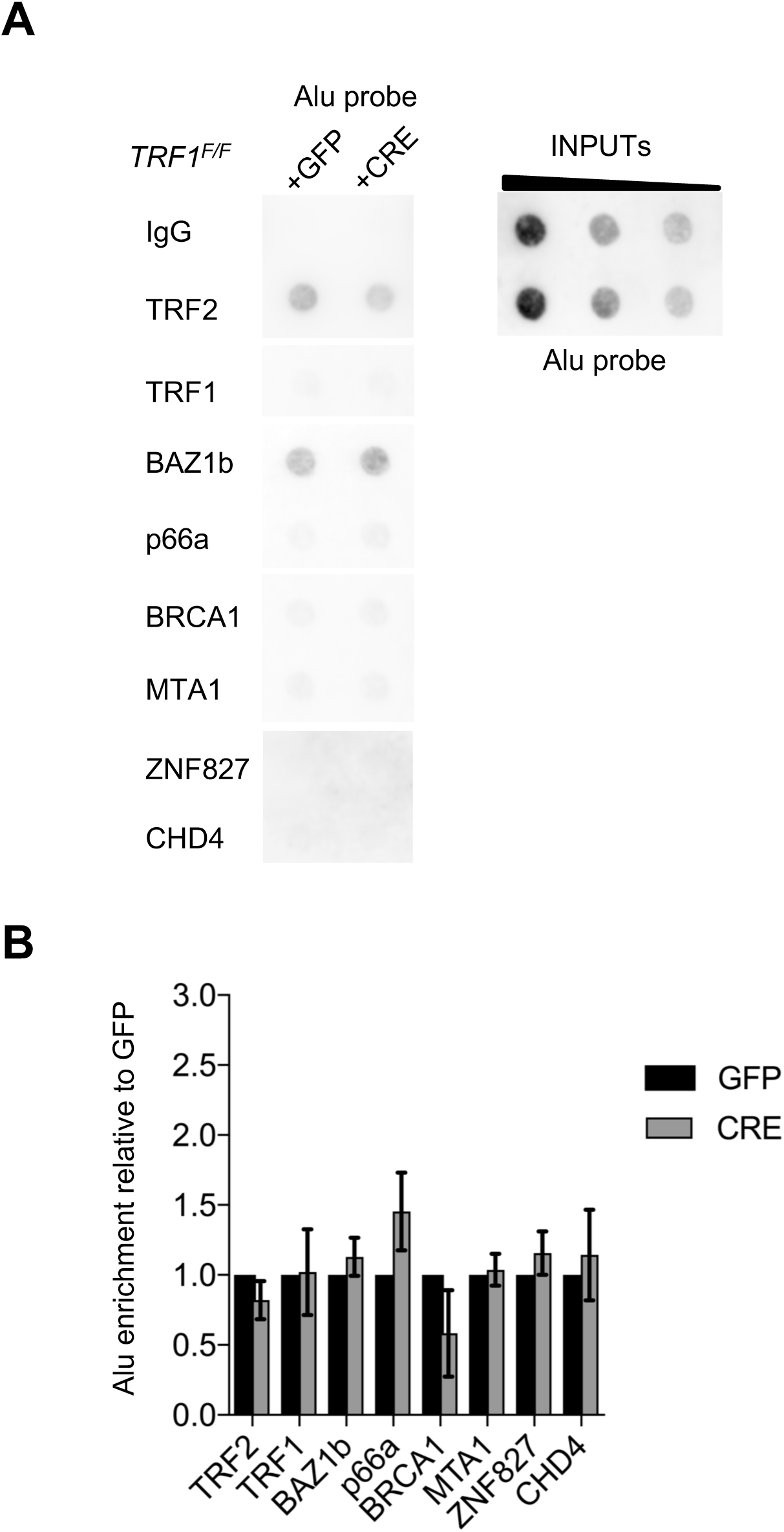
ALU control for the validation of telomeric ChIP. **(A)** Control for Figure 2B, dot-blot for validation of chromatin remodelers factor specifically recruited at TRF1 depleted telomeres. The blot was revealed with a DIG-Alu probe. **(B)** Quantification of C. ChIP signals were normalised to DNA input and GFP control and data are represented as relative Alu enrichment (n=3) ± SEM. Source data are provided as a Source Data File.

**Figure S2.**
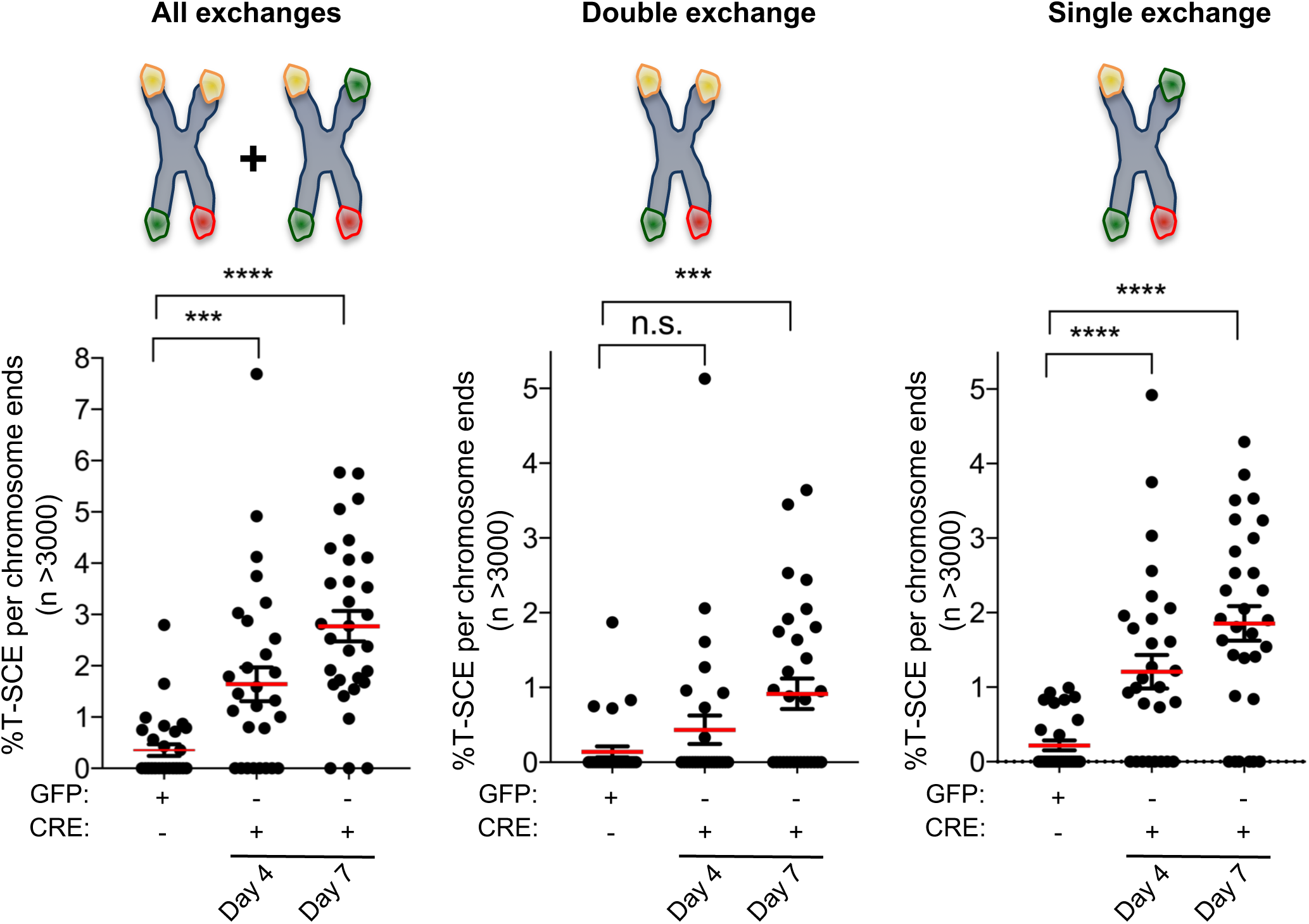
TRF1 suppresses different types of telomeric recombination. **(A)** Time course quantification of the different classes of T-SCEs using <u>denaturing</u> CO-FISH. *TRF1^F/F^* MEFs were collected 4 days and 7 days post-infection with CRE-adenovirus or GFP-(control). The different types of exchanges were classified into three different categories: all exchanges (single + double); double exchanges (reciprocal, both chromatids); single exchanges (asymmetrical, single chromatid). Graphs are representing as % of T-SCE per chromosome ends (n= at least 3000 events were scored) ± SEM. P value, two-tailed student t-test (****, *P* < 0.0001; ***, *P* < 0.001; n.s.= non-significant).

**Figure S3.**
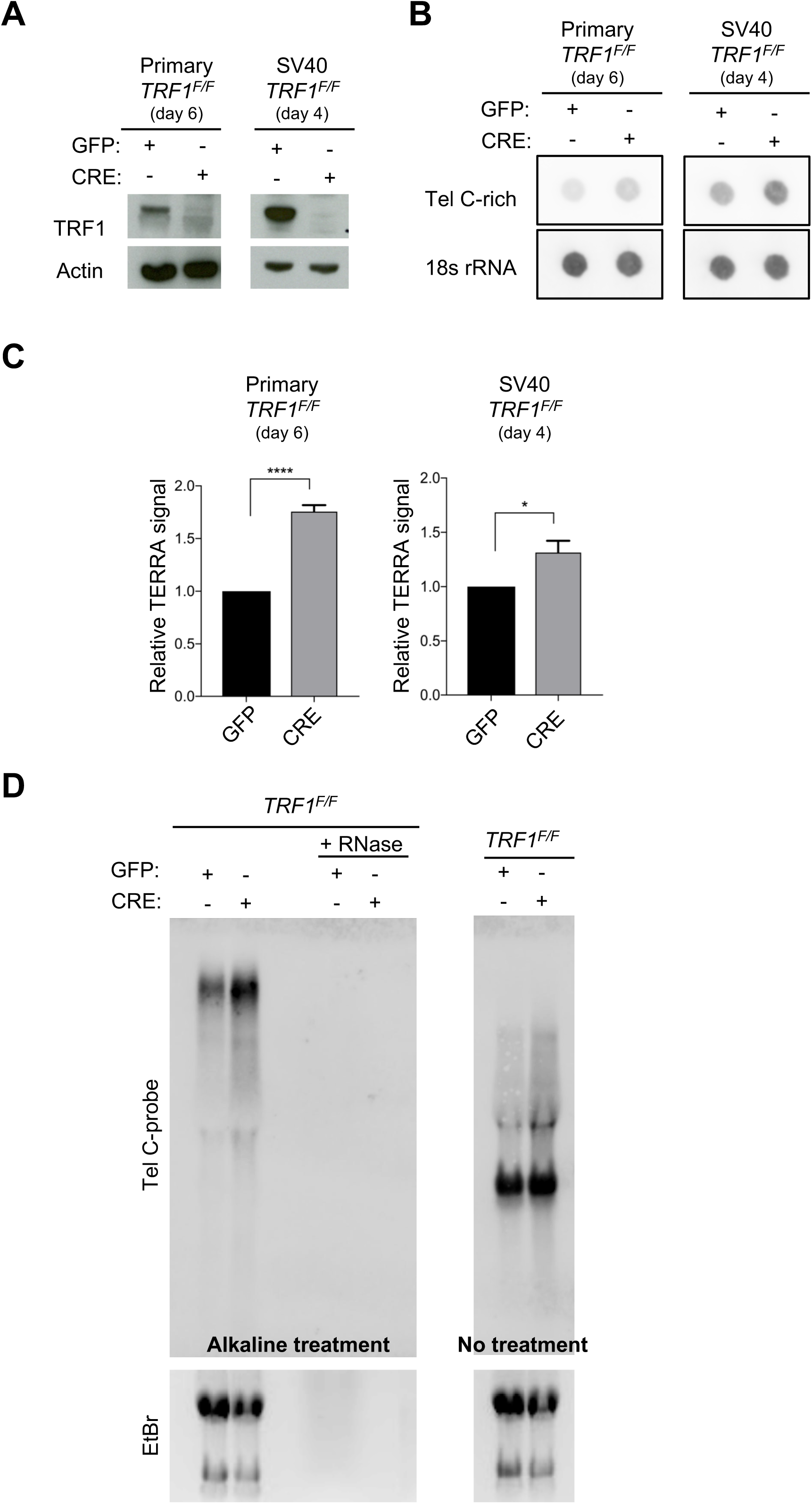
TRF1 deletion causes increased TERRA levels also in primary MEFs (day 6) and immortalised MEFs at earlier time post-infection (day4). **(A)** Western blotting showing protein expression in wt and *TRF1* deficient primary MEFs (P4) 6 days post-infection with GFP-or CRE-Adenovirus (left panel) and in SV40-immortalised MEFs, 4 days post-infection (right panel). **(B)** RNA dot-blot analysis upon *TRF1* deletion showing increased TERRA signals in CRE-infected conditions compared to control GFP-. The blot was revealed with a DIG-Tel-C-rich probe or 18s rRNA as a control. **(C)** Quantification of B. Data are shown as TERRA signal relative to GFP condition (n=3) ± SEM. P values, two-tailed student t-test (*, P < 0.05; ****, P < 0.0001) **(D)** TERRA detection by Northern blotting upon *TRF1* deletion showing increased High Molecular Weights (HMW) RNA molecules upon alkaline treatment (left blot). In native conditions, HMW-TERRAs are not detected and no significative difference is observed for low molecular weight species (right blot). The blots were revealed with a DIG-Tel-C-rich probe (upper part). Ethidium bromide (EtBr) staining (bottom) of rRNAs was used as loading control. Source data are provided as a Source Data File.

**Figure S4.**
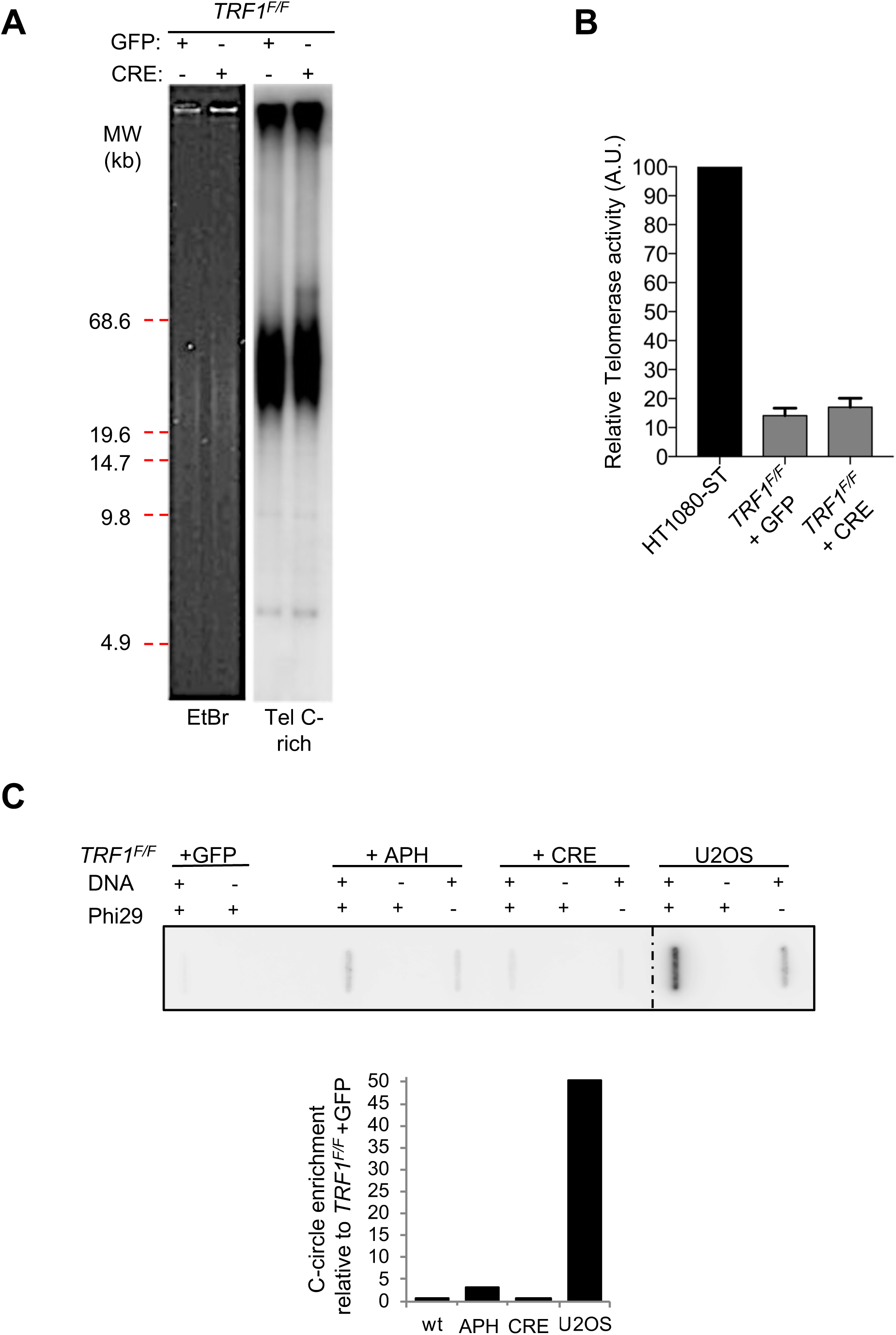
*TRF1* deficient MEFs do not present heterogeneous telomeres, neither c-circles. **(A)** Terminal Restriction Fragments (TRF) blot showing no telomere length heterogeneity upon *TRF1* deletion (+CRE, 7 days post-infection) compared to control *TRF1^F/F^* +GFP MEFs. The blot was revealed with a DIG-Tel-C-rich probe (right). Ethidium bromide (EtBr) staining (left) is used as loading control. **(B)** Quantification of telomerase activity levels by TRAP assay showing no changes in telomerase activity after *TRF1* deletion in MEFs. Values are normalised to the control HT1080-ST cells (100%) and are represented as mean (n=4) ± SD. **(C)** C-circle assay showing no c-circle formation upon *TRF1* deletion (+CRE, 7 days post-infection) compared to control *TRF1^F/F^* +GFP MEFs. Phi polymerase amplification products were spotted on the membrane and revealed using DIG-TelC probe. *TRF1^F/F^* MEFs treated with aphidicolin (APH) are used as negative control for telomere fragility not inducing C-circles, while U2OS, ALT positive cell line, is used as positive control. Source data are provided as a Source Data File.

**Figure S5.**
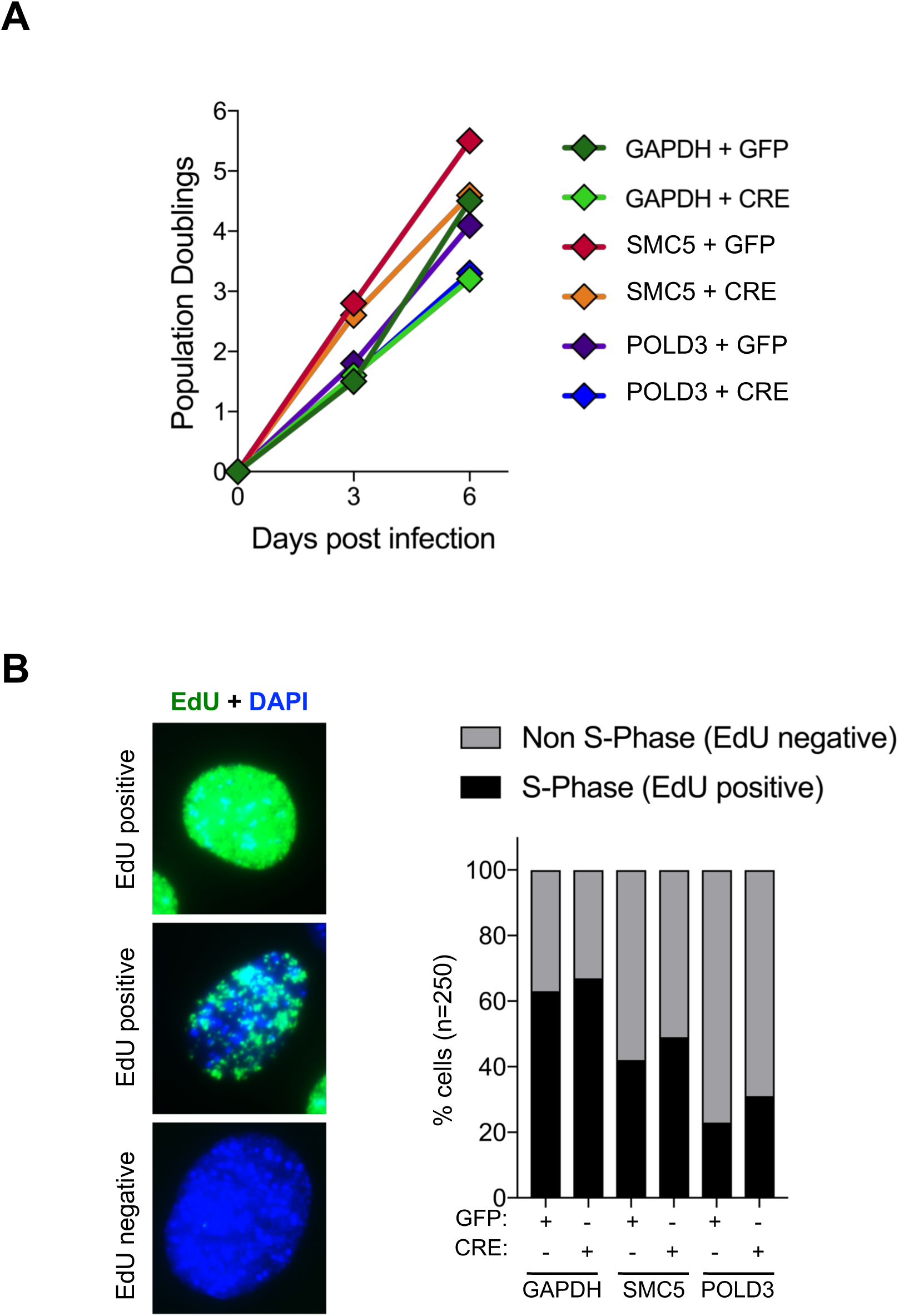
(related to Figure 4). Cell proliferation and EdU incorporation are not affected in *TRF1, TRF1-SMC5* and *TRF1-POLD3* mutants. (A) Growth curves showing cell proliferation in *TRF1^F/F^* MEFs infected with shGAPDH control (GFP or CRE), shSMC5 (GFP or CRE) and shPOLD3 (GFP or CRE). Population doublings were calculated for each condition. **(B)** Representative images of IF showing EdU(green) incorporation in MEFs nuclei (DAPI). Quantification of cells (as %) incorporating EdU using IF-staining (n=250). Cells positive for EdU staining were classified as in S-Phase, while cells negatively stained for EdU were scored as in non-S phase, for the same genetic backgrounds as in A. Source data are provided as a Source Data File.

**Figure S6.**
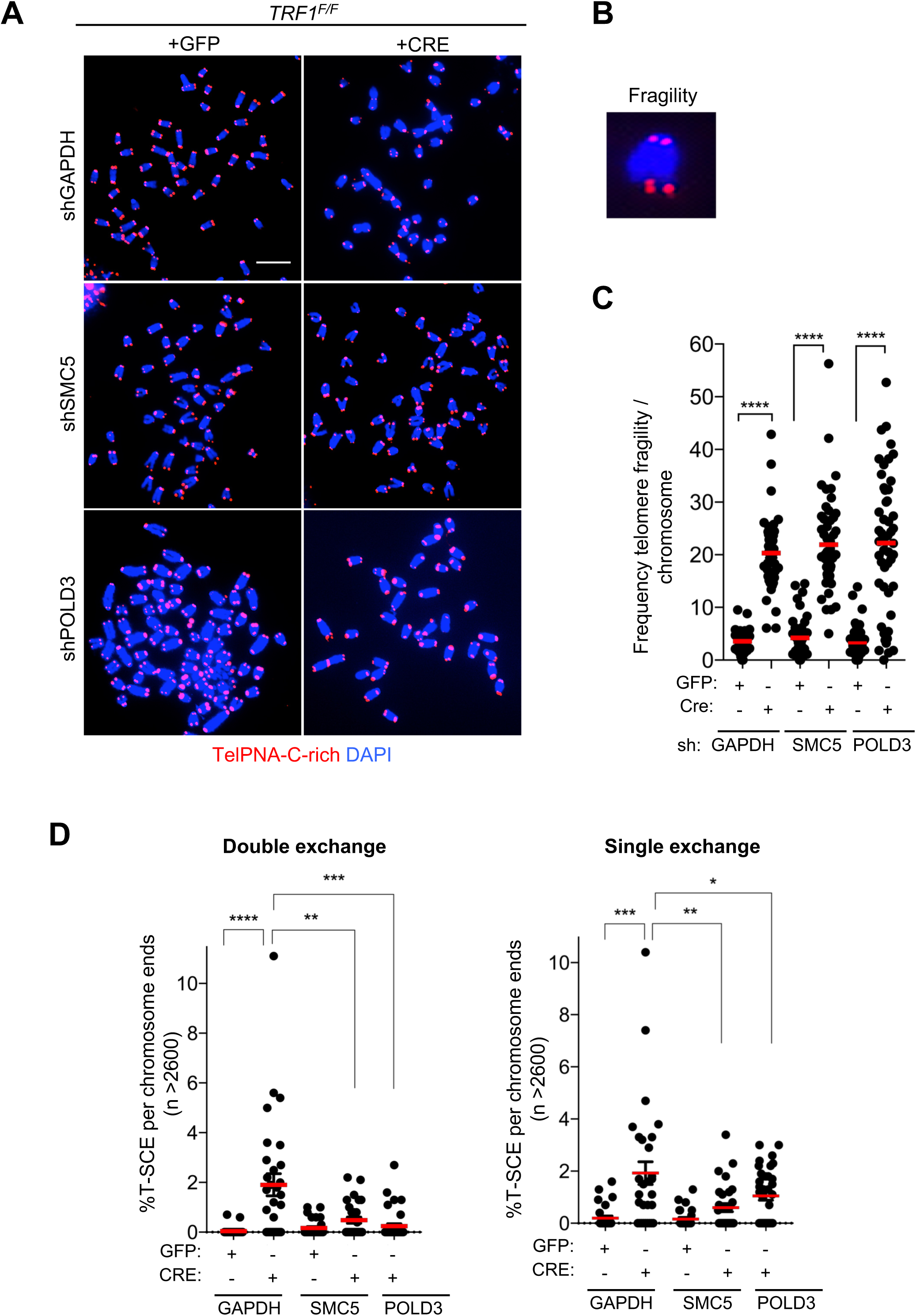
SMC5 and POLD3 are dispensable for TRF1 dependent telomere fragility but required for recombination events. **(A)** Representative images of metaphases stained with TelPNA-Cy3 probe (red) and DAPI (blue) from *TRF1^F/F^* MEFs infected with shGAPDH control (GFP or CRE), shSMC5 (GFP or CRE) and shPOLD3 (GFP or CRE). Scale bar, 10 µm. **(B)** Enlarged image showing telomere fragility. **(C)** Quantification of A-B. Data are indicated as % telomere fragility per chromosome. The mean values ± SEM are indicated. P value, two-tailed student t-test (****, *P* < 0.0001). Source data are provided as a Source Data File.

